# Commercially Relevant Species in the Mediterranean Sea: A Perspective from Late Pleistocene to the Industrial Revolution

**DOI:** 10.1101/2025.02.04.636444

**Authors:** Daniela Leal, Konstantina Agiadi, Maria Bas

**Affiliations:** University of Algarve, Campus de Gambelas, 8005-139 Faro, Portugal; Institut de Ciències del Mar (ICM-CSIC), Passeig Marítim de la Barceloneta 37-49, 08003 Barcelona, Spain; Department of Geology, University of Vienna, Josef-Holaubek-Platz 2, 1090 Vienna, Austria

**Keywords:** Fishes, Holocene, Marine mammals, Molluscs, PRISMA, Research trends

## Abstract

The Mediterranean Sea is the world’s second-largest biodiversity hotspot and has been impacted by several environmental changes and human activities since pre-historic times. We present the results of a systematic review of the published literature on the nature and extent of these impacts on the ancient–historic Mediterranean marine ecosystems. We aim to provide an overview of the current state of knowledge and identify research gaps about climate and human-activity impacts on commercially relevant species of marine mammals, fishes, and molluscs in the Mediterranean Sea over the last 130 thousand years until the Industrial Revolution (the year 1850). In most of the reviewed publications, species were used as indicators of past climatic conditions or human subsistence strategies. A research gap remains, however, in quantifying their effects on marine ecosystems. Based on our results, we identify data trends in time and space and by functional group. Data are available primarily from the Holocene rather than the Late Pleistocene, reflecting a heterogeneous availability of records. The Adriatic Sea is underrepresented among subregions, which may indicate variability of accessible data between subregions, rather than an actual lack of information. Marine mammals were less studied than fishes and molluscs in the three subregions. Despite the lack of standardised guidelines to conduct studies and subsequent variability in information, this work provides novel insights into the importance of studying the evolution of research focused on past environmental and anthropogenic impacts in the Mediterranean Sea. Research efforts need to be balanced to examine both economically and ecologically valuable species in the marine ecosystem. We also reinforce the need for uniform approaches to gather data in a usable format for posterior research.

## 1. INTRODUCTION

Marine ecosystems are extremely diverse and socioeconomically valuable globally (Millennium Ecosystem Assessment, 2005; Halpern et al., 2008). They offer cultural services (e.g., tourism) as well as ecosystem services like provision (e.g., food), regulation (e.g., water purification), and support (e.g., coastal protection) (Doney et al., 2012; Bernhardt & Leslie, 2013; Liquete et al., 2013).

The Mediterranean Sea, the world’s second-largest biodiversity hotspot (IUCN-MED, 2009), is a climate-warming hotspot (Giorgi, 2006; Diffenbaugh & Giorgi, 2012) which has been experiencing significant disruptions since prehistorical times, through climate change (e.g. Berger et al., 2016; Lionello et al., 2023) and human activities (e.g. Jackson et al., 2001; Coll et al., 2010). Environmental and anthropogenic changes significantly affect the biodiversity, functioning, and structure of the Mediterranean Sea (Lotze et al., 2006; Coll et al., 2010; Moullec et al., 2019). These impacts include habitat loss, ecosystem degradation, and the overexploitation of marine resources (Halpern et al., 2008; Coll et al., 2010; Côté et al., 2016). All these factors disrupt the natural equilibrium in marine ecosystems, affecting habitat quality, prey availability, and species phenological patterns, among others (Pauly et al., 1998; Planque et al., 2010; Howarth et al., 2014).

Despite existing knowledge about the long-term impacts of climate change and human activities on marine ecosystems (Jackson et al., 2001; Lionello et al., 2023), there is still a lack of understanding of their extent on past marine ecosystems (Cushing, 1988; Smith, 1994). Most studies focus on marine ecosystems’ current problems, often relying on data from the past century. However, these current data are already influenced by climate variations and human activities, failing to represent the pristine conditions of these ecosystems (Agiadi & Albano, 2020). This tendency can result in an underestimation of long-term changes in marine ecosystems (Pauly, 1995). To counter this trend, historical, archaeological and paleontological data are increasingly used (Lotze et al., 2006; Rick & Lockwood, 2012; Braje et al., 2017), as these records can span from hundreds to millions of years. The geohistorical record can provide long-term and pristine baseline data on marine ecosystems, offering insights beyond those of current monitoring (Dietl & Flessa, 2011). However, obtaining paleontological and archaeological data can be challenging, as it relies on the presence/absence of remains in specific locations. At the same time, historical records such as written materials, vary broadly in scale, resolution and context (Thurstan, 2022). This inherent diversity of historical sources presents a significant challenge in the construction of a comprehensive historical database.

The Mediterranean Sea is home to several emblematic and endemic species of marine mammals, fishes, and molluscs which play different ecological roles in this marine ecosystem (Beaugrand et al., 2018), from predation and grazing to ecosystem structure, functioning and engineering (Bowen, 1997; Cury et al., 2000; Wallach et al., 2015; Ysebaert et al., 2019). These marine species can influence ecosystem dynamics through their trophic position, exerting top-down, “wasp-waist” or bottom-up control on the food web (e.g., Bowen, 1997; Cury et al., 2000; Wallach et al., 2015). Filter-feeding molluscs can also improve water quality and enhance ecosystem stability (Ostroumov, 2005), while large migrators, such as marine mammals and fish, facilitate energy and nutrient transfer across ecosystems (Kitchell et al., 1979; Nathan et al., 2008).

In this context, the present work aims to assess the existing literature on climate and human impacts on commercially important species of marine mammals, fishes, and molluscs in the Mediterranean Sea over the last 130,000 years and identify research gaps. We use the Preferred Reporting Items for Systematic Reviews and Meta-Analyses (PRISMA) 2020 methodology, a widely used approach for conducting systematic reviews (Page et al., 2021). In this way, this review contributes to guiding current and future research and environmental management in the Mediterranean Sea ecosystems.

## 2. MATERIALS AND METHODS

### 2.1 Study Area

The study was carried out in the Mediterranean Sea (Figure 1), which is the largest and deepest semi-enclosed sea in the world (Robinson & Golnaraghi, 1994). It is nowadays connected to the Atlantic Ocean on the west side through the Strait of Gibraltar. To the northeast, it is connected to the Marmara Sea through the Dardanelles Strait, and further to the Black Sea through the Bosphorus Strait. In the southeast part, it is connected with the Indian Sea and the Red Sea through the Suez Strait. Physical boundaries established by the straits of Sicily (between Italy and Tunisia) and Otranto (Italy and Albania) separate the Mediterranean into three subregions: the Western and Eastern sub-basins, and the Adriatic Sea (Millot & Taupier-Letage, 2005; Rio et al., 2007; Robinson et al., 2009), which we have adopted for this work.

**Figure 1.**
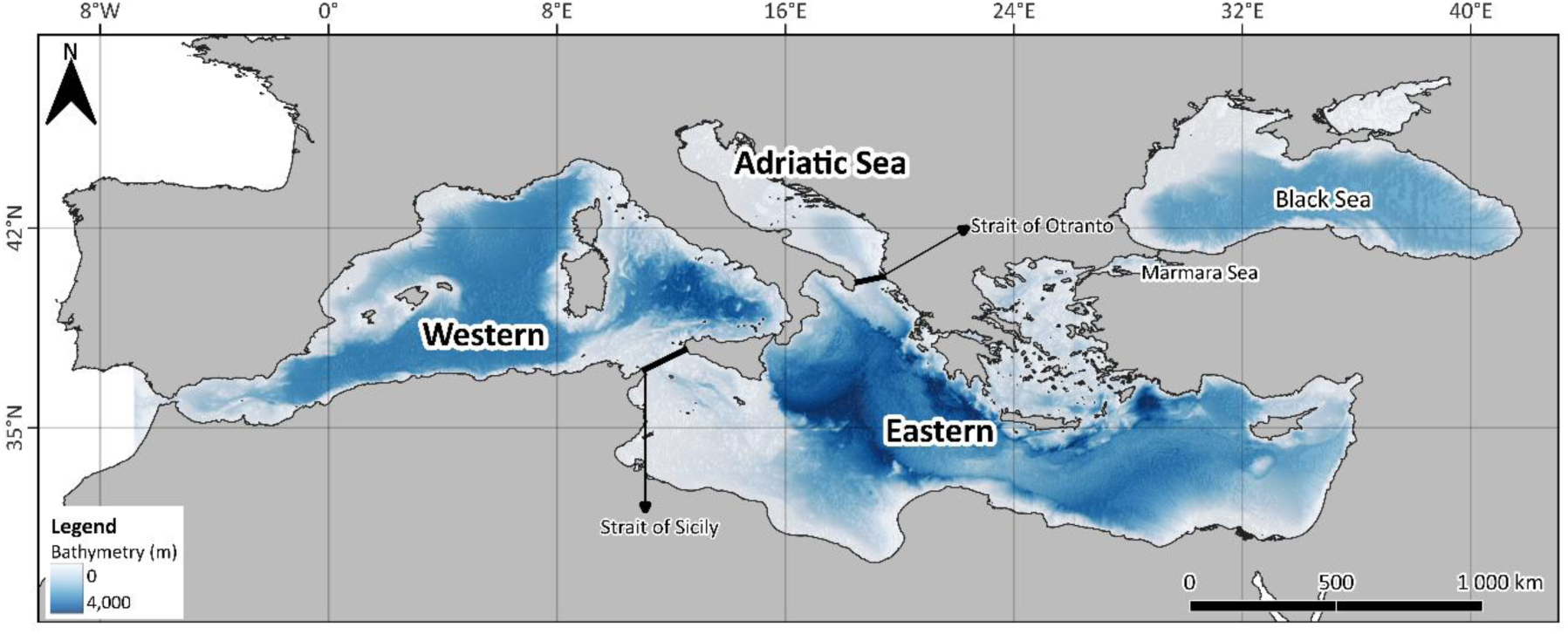
Mediterranean Sea and the regional divisions (Western, Eastern and Adriatic) adopted in this work. Black and Marmara Sea are also indicated.

### 2.2 Temporal Framework

Over the years, classification schemes for the geological, cultural, and climatic periods have been proposed for the Mediterranean Sea in different subregions. We have created a table with all the available periods so that we can perform all the analyses between these subregions and use a uniform identification of the regions and periods (see timetable from Leal et al., 2025), incorporating both the geological terms settled by the Global Boundary Stratotype Section and Points from the International Commission on Stratigraphy (ICS, https://stratigraphy.org/gssps/), as well as the cultural and environmental events terms identified in the literature. The periods in this table are dated based on years Before Present system. All conversions to this system were made using the package *rcarbon* (Crema & Bevan, 2020) in R version 4.4.1 (R Core Team, 2023).

### 2.3 Systematic Literature Review Process

The present study conducted a systematic review of peer-reviewed scientific articles, reviews, data articles and book chapters. For this work, we considered the Industrial Revolution to have started approximately around 1850 in the Mediterranean Sea (Abram et al., 2016). The Mediterranean Sea is characterised by high biodiversity and a long history of interaction between humans and marine resources (Coll et al., 2010 with references therein). This is why we decided to focus on commercially important species (including marine mammals, fish and molluscs) most commonly found in records referring to the use, consumption and/or exploitation during the past (see Table 1 for all the species used in this work). This work was conducted using the PRISMA 2020 methodology (Preferred Reporting Items for Systematic Reviews and Meta-Analyses; Page et al., 2021). This methodology is divided into four steps: (1) a systematic selection of peer-reviewed articles using search engines; (2) removal of duplicates and two screening processes based on pre-set inclusion/exclusion criteria, (3) identification of other relevant studies via secondary literature, and (4) the extraction of information from both sources (search engines and secondary literature) into a database based on targeted variables. As previously mentioned, The final search included a total of seventeen species: two marine mammals, twelve fishes, and three molluscs.

**Table 1.**
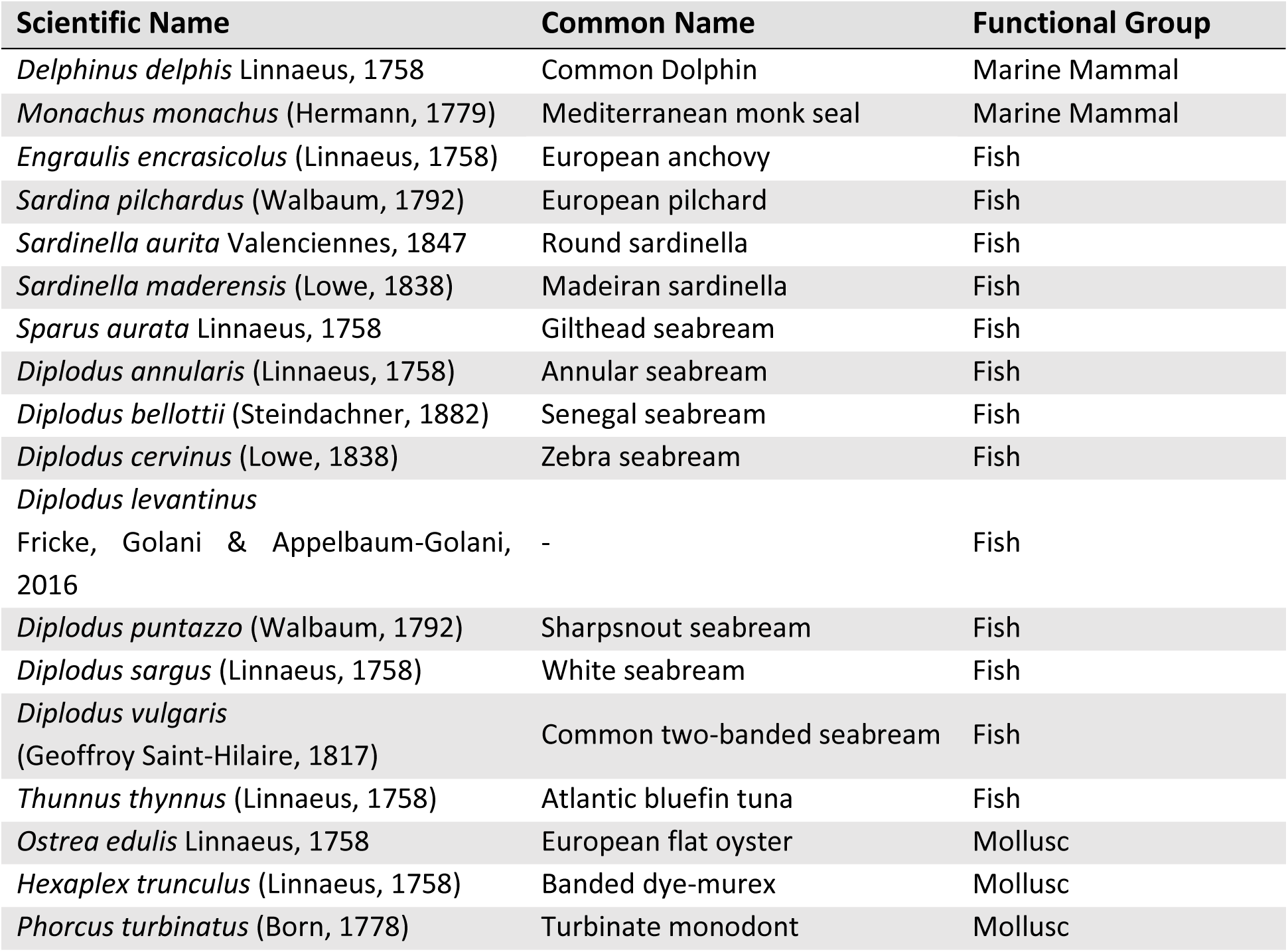
Species analysed in this work.

The searches were conducted in English. The databases used to retrieve the information were Elsevier’s Scopus (https://www.scopus.com/) and the Web of Science Core Collection (https://webofknowledge.com/). The search covered the interval from 1911 until the cut-off date, February 12, 2024. We searched using the “All fields” mode, which explores all the searchable fields using one query, including “title”, “abstract”, and “keywords”. The strings used in the search were grouped into five main categories with a specific set of keywords within – taxa, with currently scientifically accepted names for each species, reported by the World Register of Marine Species (WoRMS; https://www.marinespecies.org), the most commonly used unaccepted names found in literature and their respective common names; subregions of interest; timeframe; impacts including both human and climate; and theme. The keywords from each topic were combined using the Boolean operator “OR”, while different topics were connected using the operator “AND”. The “impact-related” topic terms were combined in a single query to avoid restrictive results (see Table S1 to check all strings used within each category). Once the searches were done in both search engines, the datasets were merged, the duplicates were removed, and the remaining articles were screened.

The screening was divided into two steps. In the first step, the titles and abstracts were screened considering the targeted species, study area and timeframe. Only the articles meeting the first inclusion criteria (Table 2) were downloaded and used for the second screening. In the second screening, only the papers with quantitative data, which also met the inclusion criteria (Table 2) based on the full-text screening, were included in the database. See Table S2 for the compilation of scientific publications used to build the database. With this information, a database was built (Leal et al., 2025), including a set of variables that captured the details of the included article. These variables encompassed general information about the articles, the location of the data (coordinates), the type of assemblage from where the data originated (categorised as natural or archaeological), and the respective age (absolute or relative). In cases where the authors did not provide specific coordinates but indicated the general of the site, we assigned the coordinates representing the midpoint of the area. Additionally, we collected information on certain effect measures, such as the dating method used, the time interval covered by the study and the sampling effort.

**Table 2.**
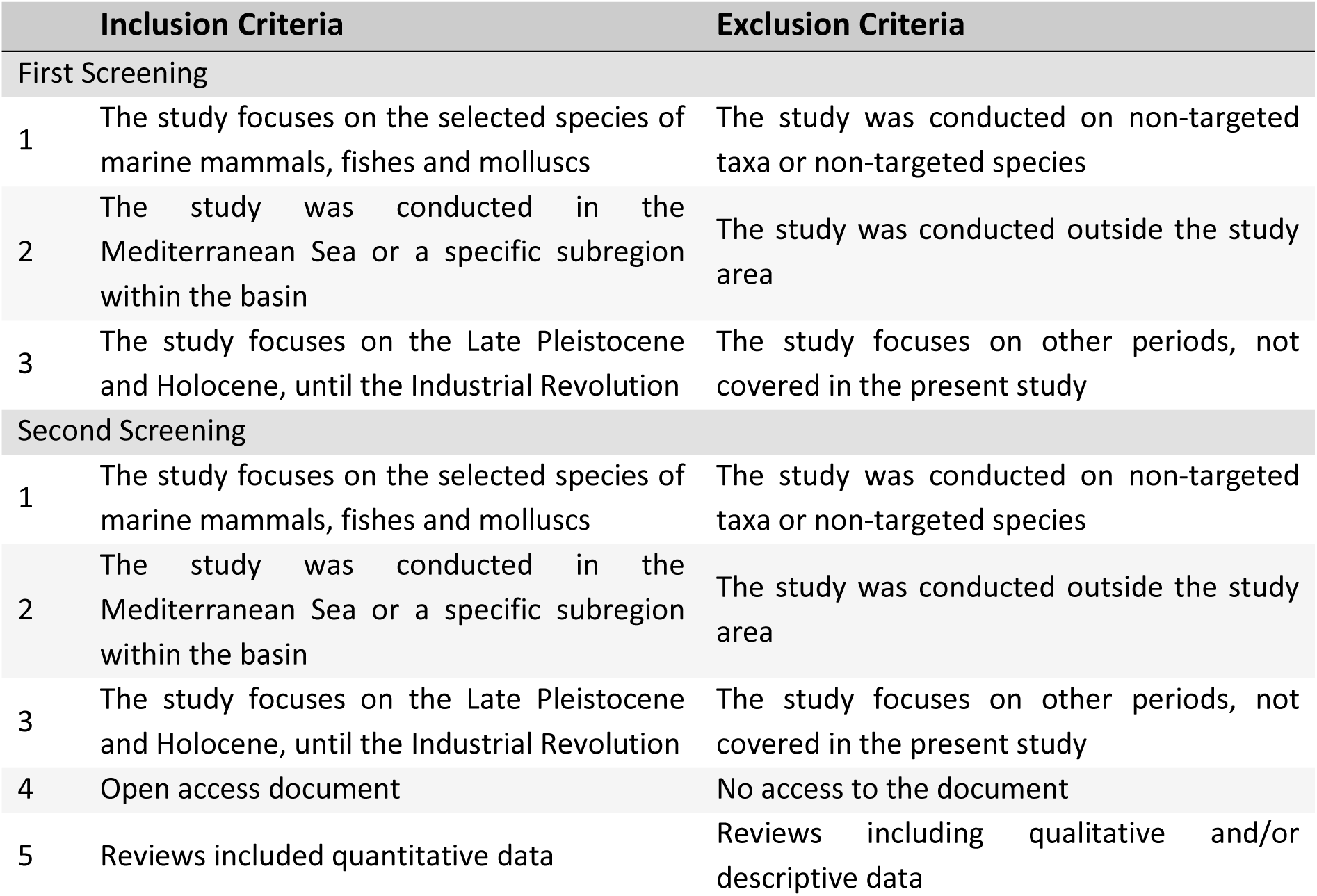
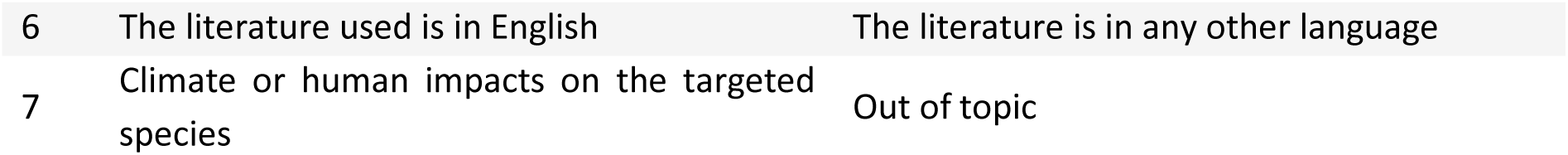
Inclusion and exclusion criteria for literature screening in the systematic review.

For other specific variables used for subsequent quantitative analyses, we chose not to explore this topic in the present work, as it falls outside the scope of the general objective outlined above.

### 2.4 Data Analyses

All the maps presented in this work were created using Free and Open Source QGIS version 3.34.3 (QGIS.org). The bathymetry data was obtained from GEBCO (General Bathymetric Chart of the Oceans) (GEBCO Compilation Group, 2022). All original graphical representations were made with the *ggplot2* package (Wickham, 2016) and statistical analyses were conducted in R version 4.4.1 (R Core Team, 2023).

To analyse publication trends over time, we categorised the data by marine fauna and then by functional group. Firstly, we plotted the cumulative number of publications per year for the entire Mediterranean Sea and subregion. Secondly, we examined publication trends for each functional group, applying the same method of plotting the cumulative number of publications per year for the Mediterranean Sea and each of the three subregions.

The data on effort measures were intended to help readers understand the level of effort the authors invested in researching these particular species across the Mediterranean Sea. The variables used included the number of study sites, the age certainty method and the temporal coverage provided by each study. When the authors provided the temporal coverage as relative age, such as cultural periods across the three subregions, the absolute age information was complemented with data from the timetable (Leal et al., 2025). For the dating system variable used in the publications, we focused on identifying the most commonly applied methods in the dataset. We also calculated central tendency and variability statistics for the remaining variables along with the distribution of publications per geological period, which provides an overview of the most studied periods, as shown in Table 3.

**Table 3.** Temporal coverage of each geological period. ICS refers to the International Commission on Stratigraphy (Cohen et al., 2013).

| Years BP | Series | Stage [ICS] | Reference |
| --- | --- | --- | --- |
| 4,200 - 0 | Holocene | Late Holocene | Stuiver and Reimer (1986) |
| 8,186 – 4,200 | Holocene | Middle Holocene | Stuiver and Reimer (1986) |
| 11,650 – 8,186 | Holocene | Early Holocene | Stuiver and Reimer (1986) |
| 131,949 - 11,650 | Pleistocene | Late Pleistocene | Head et al. (2021) |

To analyse the distribution of the targeted species across subregions of the Mediterranean Sea within the defined timeframe, we first grouped the same sampling sites into sampling units. Different articles presented the same sampling location, thus, to offer a clearer overview of the main data collection sites across the Mediterranean Sea we decided to merge the same sampling sites into unique sampling units (Table S3). Additionally, sampling sites located in both the Black and Marmara Seas were included in the Eastern Mediterranean due to the limited number of sites in these seas and to standardise the number of regions within the Mediterranean basin.

To analyse the distribution of the selected species in time, we visualised the records of marine mammals, fishes, and molluscs across four geological periods (from the Late Pleistocene to the Late Holocene) using pie charts overlaid on a map (see Table 3 for the temporal coverage of each period).

## 3. RESULTS

### 3.1 Systematic Literature Review

A total of 2,762 publications were retrieved from both Web of Science and Scopus. After removing duplicates, 1,523 publications were screened (Figure 2). From the first screening, 117 publications were considered eligible for full-text screening. From the second screening, 40 publications were included for gathering information and subsequent analyses. We also included 8 additional publications in the analyses through secondary literature. In total, 48 publications were used to build the database (Figure 2). The publications were also categorised based on their primary focus, either related to the species as indicators of past environmental conditions or related to previously identified climate and/or anthropogenic impacts upon them. With this classification, 42 of the total number of publications analysed targeted species as indicators and 15 assessed specific climate and human impacts upon the species.

**Figure 2.**
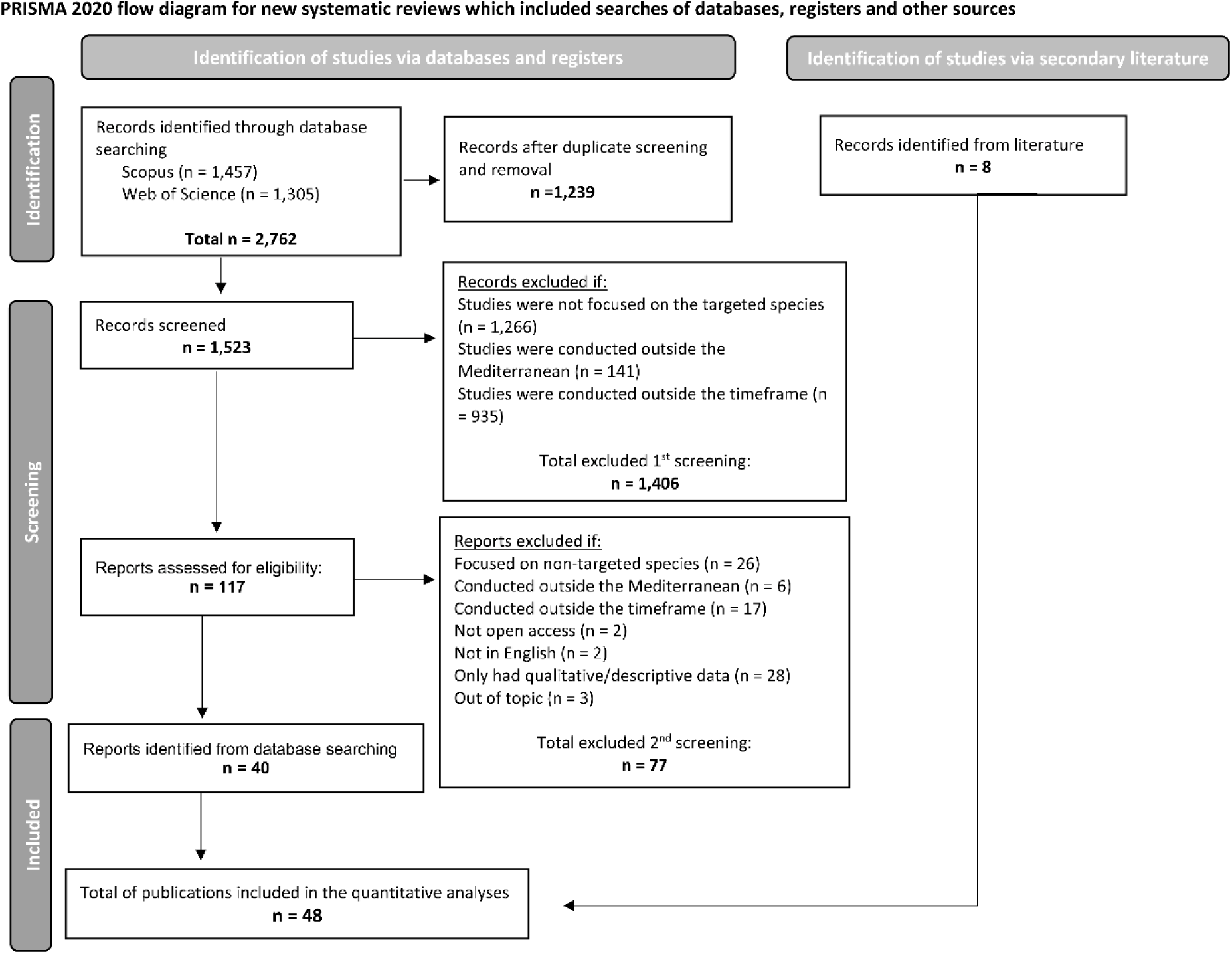
PRISMA 2020 flow diagram adapted from Page et al., 2021, illustrating the methodology and selection process used, indicating the number (n) of studies included in this review. Bold numbers indicate key values related to the number of studies included in the evaluation process.

The search covered seventeen species, but the results did not represent them all equally. No results were found for the Senegal seabream, Zebra seabream, *Diplodus levantinus*, and the Sharpsnout seabream. Instead, the search yielded data on twenty-three taxa: Clupeidae, Delphinidae, Phocidae, Sparidae, and Scombridae families. *Diplodus spp., Thunnus spp., Phorcus spp.,* and *Monodonta sp*. genus. Common dolphin, Mediterranean monk seal, European anchovy, European pilchard, round sardinella, Madeiran sardinella, gilthead seabream, annular seabream, white seabream, common two-banded seabream, Atlantic bluefin tuna, European flat oyster, banded-dye murex and turbinate monodont.

### 3.2 Research Trends

The distribution of publications over the years appeared to follow an increasing trend, with a more significant increase starting from 2015 onwards (Figure 3). When comparing the three subregions, publications concerning the Western and Eastern Mediterranean have been available over the years, while publications for the Adriatic Sea began around the year 2010. The publication trend for the Western Mediterranean seemed to follow an increasing pattern, aligning with the general trend. In contrast, the Eastern Mediterranean showed an increasing trend as well, but between 2010 and 2016, there was a slight decrease, followed by a high increase. For the Adriatic Sea, the publication trend appeared relatively constant over the years (Figure 3).

**Figure 3.**
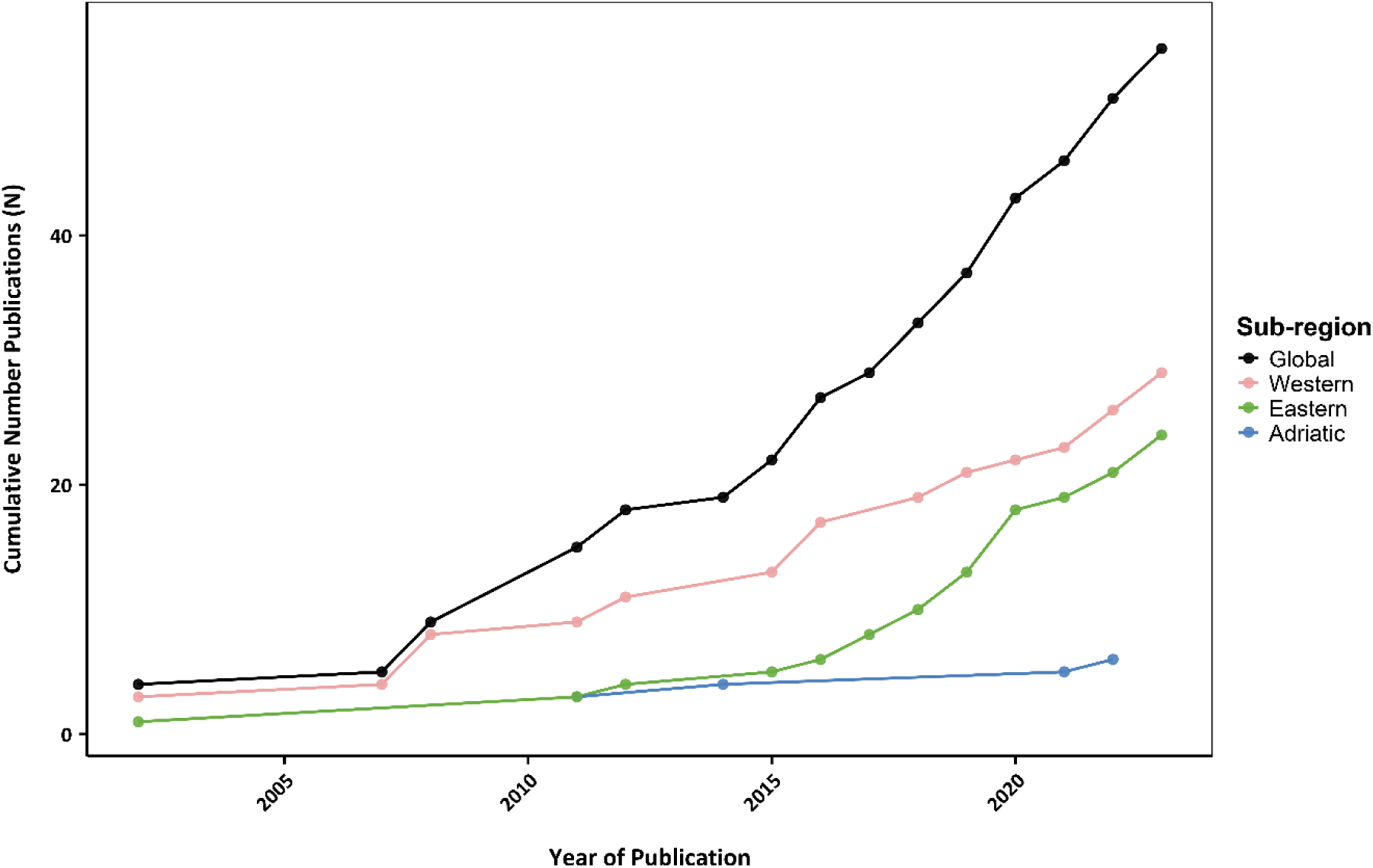
Cumulative number of publications through time for the studied marine fauna, categorised by data for the entire Mediterranean Sea (black) and each subregion (coloured).

When comparing publication trends among the functional groups, there were differences (Figure 4). Publications on marine mammals were fewer compared to those on fish and molluscs. For marine mammals, the overall trend was positive and increasing, with the Western and Eastern Mediterranean showing similar trends, although the Eastern Mediterranean had fewer publications. For the Adriatic Sea, no clear trend was established due to only one publication (Figure 4).

**Figure 4.**
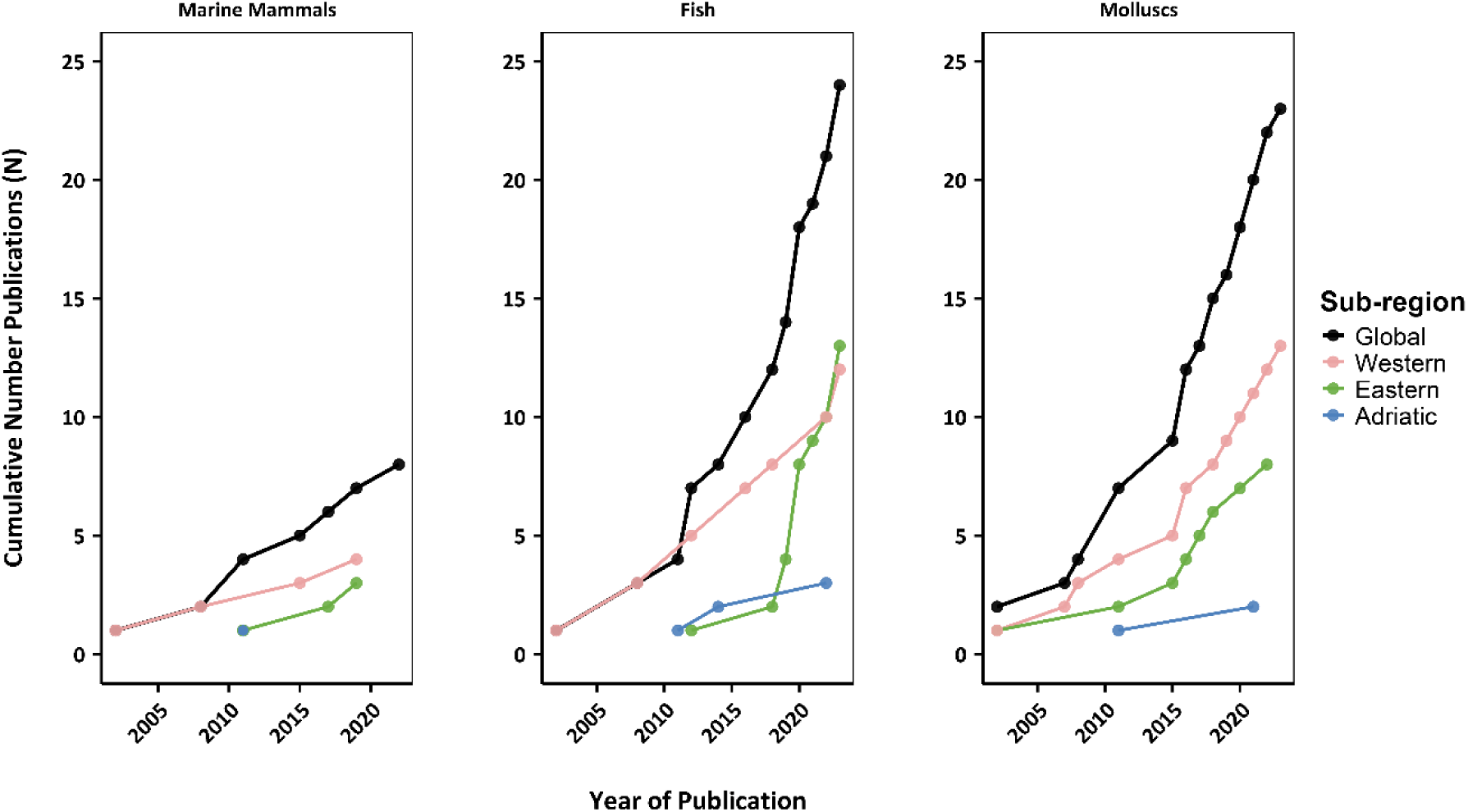
Cumulative number of publications through time, for marine mammals, fish and molluscs, categorised by data for the entire Mediterranean Sea (black) and each subregion (coloured).

For fish, the general trend showed a steep increase over time, though the trends differ among subregions (Figure 4). The Western Mediterranean displayed a positive but less pronounced trend than the overall trend. The Eastern Mediterranean had fewer publications in the initial years, followed by an increase from 2017 to the present. Research on the Adriatic Sea followed a positive trend but remained more constant over time (Figure 4).

Molluscs showed a generally positive publication trend over time (Figure 4). Both the Western and Eastern Mediterranean exhibited positive trends, although the Eastern Mediterranean experienced a decrease between 2010 and 2015. The Adriatic Sea also showed a positive trend, though it remained more stable than the other subregions (Figure 4).

The number of sampling sites per study ranged from 1 to 270, with a mean value of 33 sampling sites (Table 4). In terms of temporal coverage, the mean coverage (∼241 million years) is significantly higher than the median (∼5 thousand years), ranging between 84 and 100.5 million years (Table 4). The number of publications per geological period showed that the Late Holocene had the most studies, in contrast to the other periods. The results also showed that radiometric dating was the most used method (n=23; Leal et al., 2025).

**Table 4.** Summary of descriptive statistics for effort measures.

| Number of sites per study |  | Temporal coverage (years) |  | Number of publications per geologic period |  |
| --- | --- | --- | --- | --- | --- |
| Mean | 33.3 | Mean | 241,3101 | Late Pleistocene | 14 |
| Median | 10 | Median | 5295.5 | Early Holocene | 15 |
| Minimum | 1 | Minimum | 84 | Middle Holocene | 15 |
| Maximum | 270 | Maximum | 100.5 Ma | Late Holocene | 27 |

### 3.3 Spatial Distribution of the Data

The sampling units were not equally spatial distributed throughout the Mediterranean area (Figure 5). The Western and Eastern Mediterranean had a higher number of sampling units containing the targeted species compared to the Adriatic Sea. Additionally, isolated sampling units were presented in the Black and Marmara Seas (Figure 5, see also Table S3 for detailed information on each sampling unit). Additionally, most of the sampling units were located in the coastal areas.

**Figure 5.**
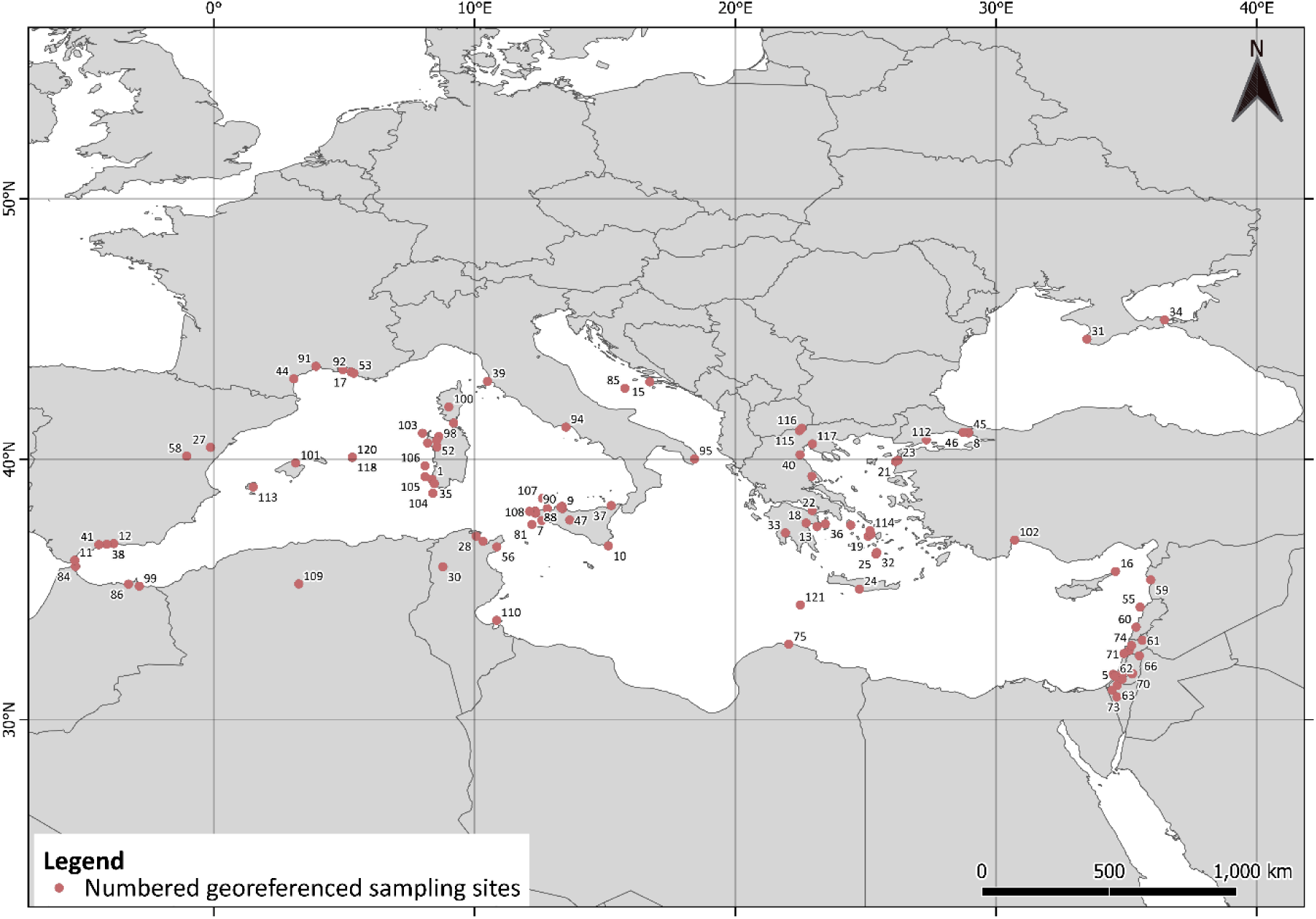
Map of the Mediterranean Sea showing sampling units (red circles). Each number corresponds to a unique sampling site. Detailed information on each location and its corresponding publication is provided in Table S3.

### 3.4 Functional differences through time

There were differences when comparing the distribution of marine mammal records across the three subregions (Figure 6). In the Western Mediterranean, there was a higher percentage of records from the Late Pleistocene period, particularly from the Phocidae family. Conversely, the Early Holocene period exhibited a similar percentage of records between Phocidae, Delphinidae, and the common dolphin. In the Adriatic Sea, records were distributed across the three periods, with a lack of data for the Early Holocene. It is important to note the shift from a higher percentage of records from the Delphinidae and Phocidae families to a predominance of common dolphin records in more recent years. In the Eastern Mediterranean, the two most recent geological periods showed a similar percentage of common dolphin records. Interestingly, the Late Pleistocene period included only records of the Mediterranean monk seal, whereas the Early Holocene period exhibited records from both families.

**Figure 6.**
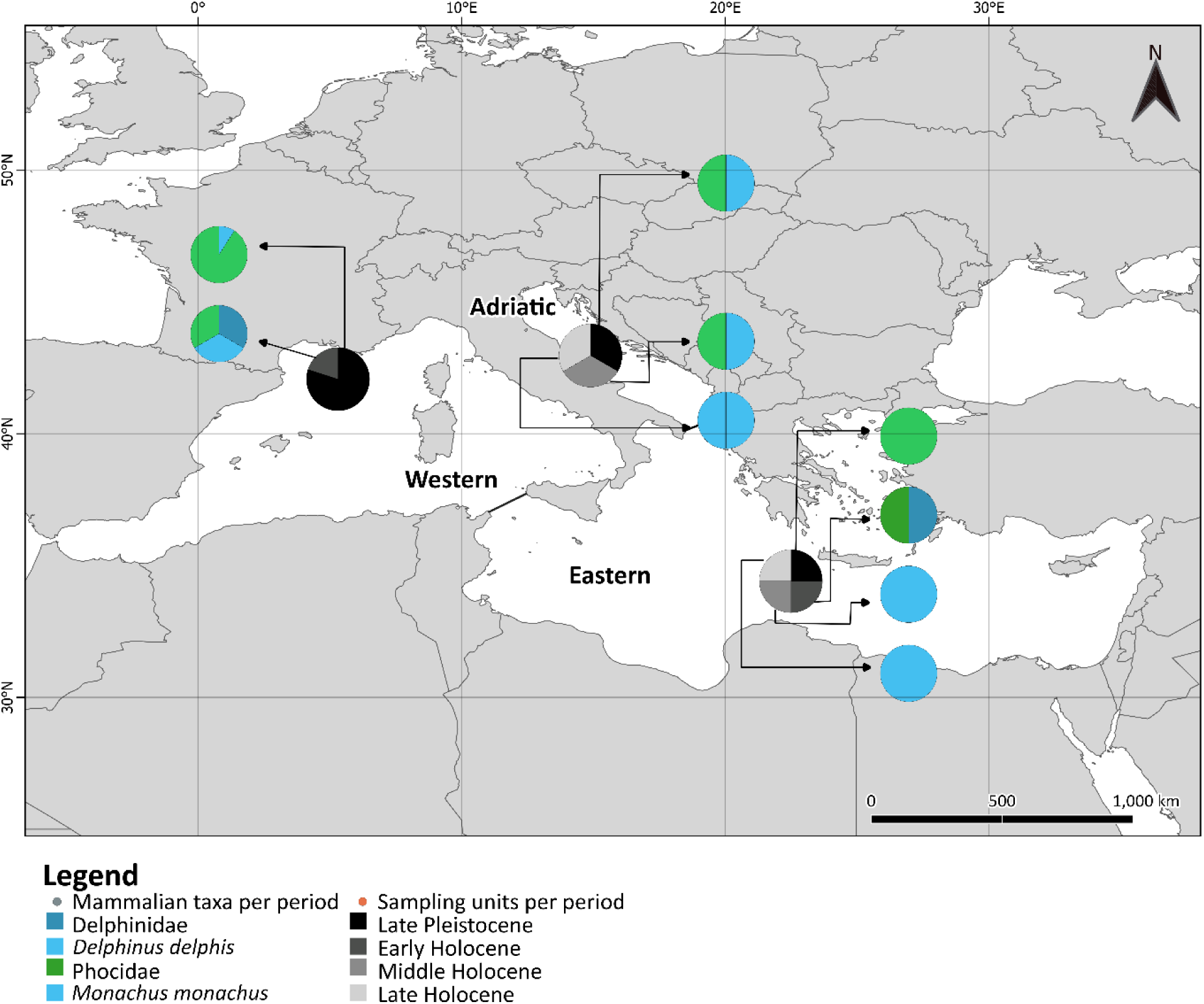
Proportional distribution (%) of sampling units per geological period, species and corresponding family across Mediterranean subregions for marine mammal records. The central pie chart (black to light grey) represents the proportion of records per geological period, while secondary pie charts show the functional distribution of records for each period. Taxa within the same family are represented using gradient colours.

The distribution of fish records was consistent across the studied Mediterranean subregions (Figure 7), though variability existed within the periods for each subregion. For example, there was a lack of data for the Late Pleistocene in the Eastern Mediterranean, while in the Western Mediterranean, records were concentrated in the Late Holocene. Atlantic bluefin tuna and Sparidae records dominated across all periods, with the presence of the Clupeidae and Scombridae families. In the Adriatic Sea, the Middle Holocene showed the highest concentration of data, with Clupeidae being the most common family across all periods, complemented by significant contributions from the gilthead seabream and the Atlantic bluefin tuna. The Eastern Mediterranean again lacked information for the Late Pleistocene. When examining the entire Holocene and all species, the Atlantic bluefin tuna consistently presented a higher percentage of records. Notably, fish diversity increased throughout the Holocene, reflecting a rise in the variety of recorded species.

**Figure 7.**
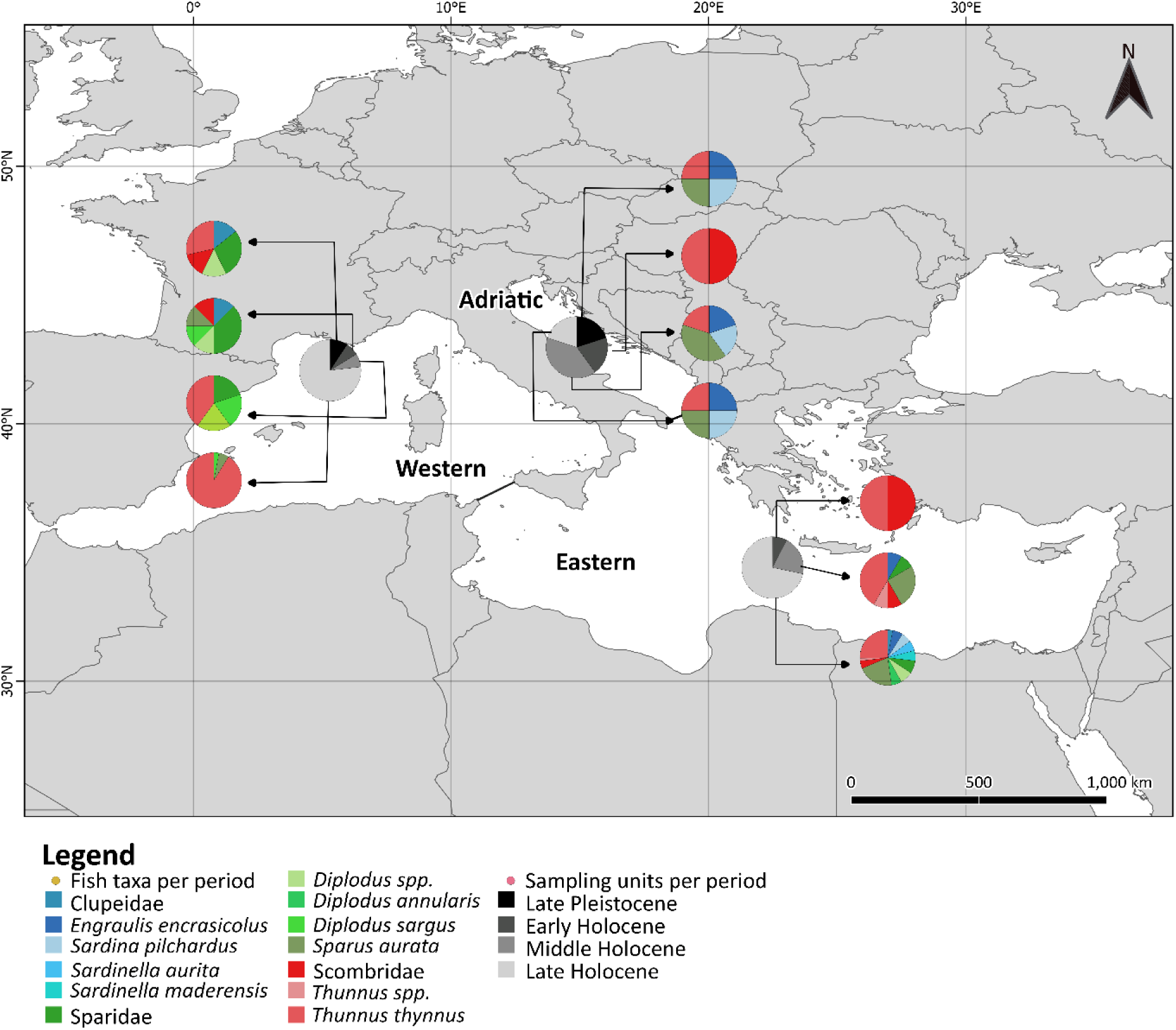
Proportional distribution (%) of sampling units per geological period and corresponding taxa for fishes by each subregion of the Mediterranean Sea. The central pie chart (black to light grey) represents the proportion of records per geological period, while secondary pie charts show the functional distribution of records for each period. Taxa within the same family are represented using gradient colours.

There were mollusc data available for all subregions and all geological periods (Figure 8), although in varying percentages. The Western Mediterranean had more records for the Early Holocene, the Adriatic Sea for the Middle Holocene, and the Eastern Mediterranean from the Late Pleistocene and Late Holocene. Turbinate monodont records accounted for a high percentage across the three subregions and four geological periods. The European flat oyster showed the highest percentage in the Western Mediterranean, while the banded-dye murex dominated in the Eastern Mediterranean. In the Adriatic Sea, both species were present, but turbinate monodont consistently had the highest percentage across all periods.

**Figure 8.**
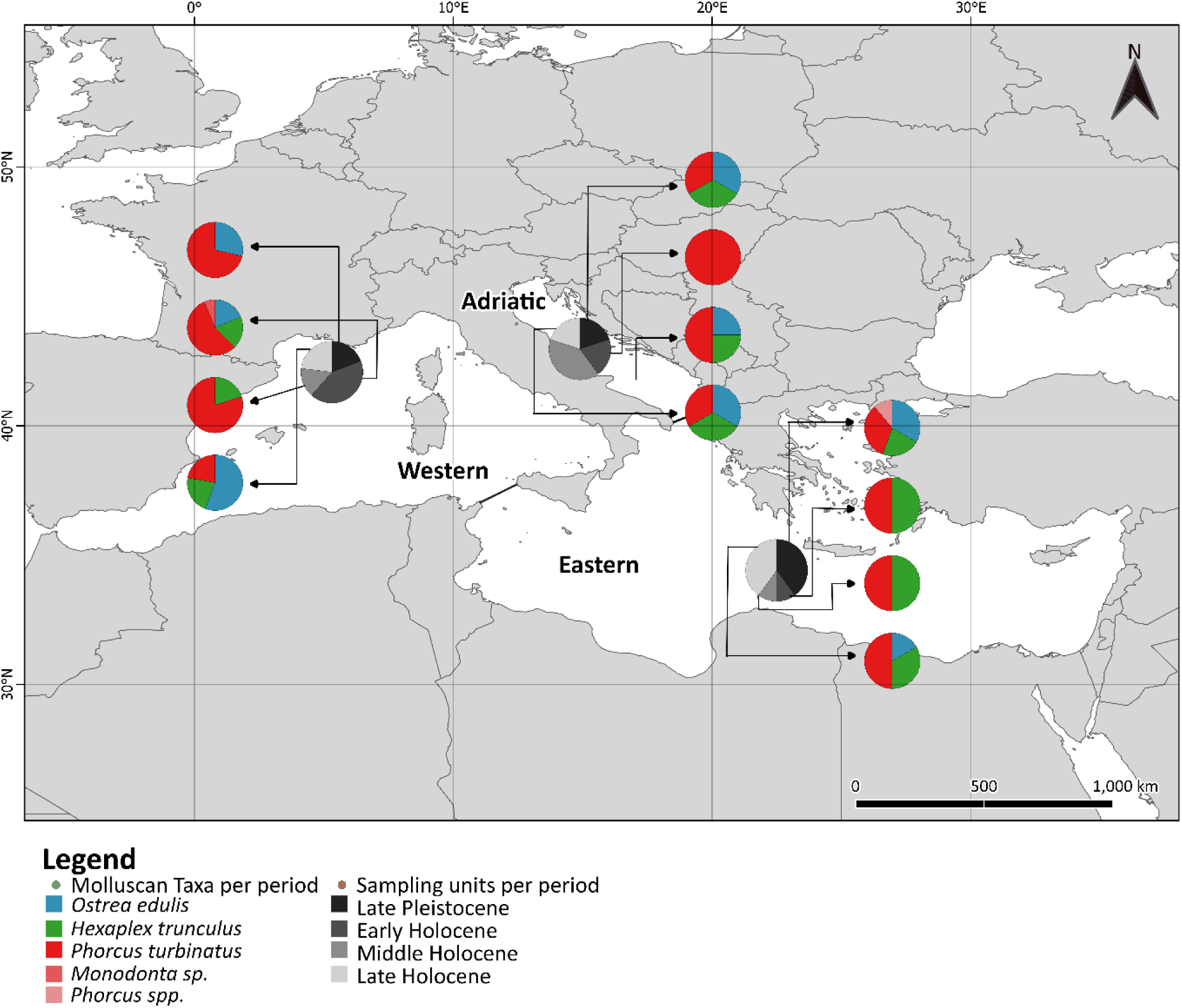
Proportional distribution (%) of sampling units per geological period and corresponding taxa by each subregion of the Mediterranean Sea for molluscs. The central pie chart (black to light grey) represents the proportion of records per geological period, while secondary pie charts show the functional distribution of records for each period. Taxa within the same family are represented using gradient colours.

## 4. DISCUSSION

Our systematic review reveals an increasing temporal trend in research focused on marine mammals, fishes and molluscs that may have been affected by climate and anthropogenic pressures over the last 130,000 years. The Adriatic Sea seems to be underrepresented across all functional groups, with marine mammals being particularly less studied than fishes and molluscs. The data primarily originate from the Holocene, reflecting the availability of written and archaeological records, compared to Late Pleistocene information.

Most publications in the database focus on multiple species, using them as indicators of paleoclimatic and paleoenvironmental conditions or human subsistence strategies (e.g., Bosch et al., 2018; Branscombe et al., 2020; Segaoui et al., 2022). This highlights a potential research gap: while long-term data exist to evaluate past climate and human impacts, relatively few studies address these effects on marine ecosystems. For example, species like the turbinate monodont, which is highly common in archaeological deposits in the Mediterranean Sea (Mannino et al., 2007, 2008), are often used for reconstructing past conditions rather than assessing the potential impact on this species and the ecosystem. Therefore, there is a need to apply a more inclusive approach in targeting multiple species that could enhance our understanding of historical ecosystem changes (Jackson et al., 2001; Lotze et al., 2011; Thurstan, 2022).

The increasing number of publications since 2015 across the Mediterranean Sea suggests a growing research interest in past impacts. However, regional imbalance persists with more studies conducted in the Western and Eastern subregions compared to the Adriatic Sea. This discrepancy might reflect the availability of accessible data rather than an actual lack of information about the Adriatic Sea. Studies conducted in this subregion such as Lotze et al. (2006) highlight the current research, with important implications regarding ecosystem-level changes. However, despite the existence of this type of research, there is a need for more targeted data collection in underrepresented subregions to assess the potential impact on the Mediterranean ecosystem. Temporal analyses show a predominance of Holocene records over Late Pleistocene data. This disparity is likely due to better-preserved, more accessible Holocene records (e.g., Andrews et al., 2023; López-Sáez et al., 2023). While some studies span extensive timeframes (e.g., Andrews et al., 2022; Lotze et al., 2011), others focus on shorter periods, further emphasising the need for comprehensive, standardised data collection.

This systematic review showed considerable variability in the research effort metrics. Differences in the number of sites and temporal coverage per study depend on the sampling design and the contents of each sampling location, lacking equality in data provided by each publication. Although several dating methods were employed across publications, the most used absolute dating method in the database is radiometric dating (e.g., Bosch et al., 2018; Colonese et al., 2018; Agiadi & Albano, 2020). This method attributes age to the decay of the natural radioactive components (Wiens, 2002), which offers higher accuracy than other dating methods, such as contextualisation of the stratigraphic layers/units or archaeological context.

Regarding the research focused on each functional group, marine mammals seem to be the least represented in spatial and temporal records compared to fish and molluscs. Despite their ecological and cultural significance, marine mammals face ongoing challenges from overexploitation, bycatch, and pollution (Schipper et al., 2008). Identifying their remains in archaeological contexts is particularly difficult due to fragmentary evidence (Mulville, 2002), but advances in genetic methods (e.g., ancient DNA extraction; Lindqvist et al., 2009) offer promising solutions. Greater use of such techniques could significantly improve data availability for this group. Iconic species like the Mediterranean monk seal and common dolphin were historically abundant but have experienced steep declines due to hunting and habitat destruction (Johnson, 2004; Lotze et al., 2011). Similarly, commercially valuable species like the Atlantic bluefin tuna and Sparidae family show extensive exploitation histories, with some species continuing to support modern fisheries (e.g., García-Vargas & Florido del Corral, 2010; Di Natale, 2014). Molluscs, particularly the turbinate monodont, are well-documented in archaeological records, often reflecting their ecological resilience and ease of harvesting (e.g., Bosch et al., 2018; Lo Presti et al., 2019).

Although this work provides novel information on how the research focused on past impacts in the Mediterranean Sea has been addressed scientifically and its research gaps, it has some limitations. These include the absence of standardised sampling methodologies across studies (e.g., absolute dating method and respective dating systems), the diversity of publication types and the variability in data. The data availability across functional groups may be linked to their commercial value. While emblematic species such as cetaceans and pinnipeds remain underrepresented (e.g., Mulville, 2002; Tortosa et al., 2002), species that are of both commercial and ecological importance, including Atlantic bluefin tuna (e.g., García-Vargas & Florido del Corral, 2010; Di Natale, 2014; Andrews et al., 2022) and Sparidae (e.g., Basurco et al., 2011; Colonese et al., 2018; Guy et al., 2018) receive more research focus. To overcome these constraints, we reinforce the necessity for balanced research efforts to be made, to study species that play significant ecological roles alongside those of economic interest. In addition, further efforts should be made to establish structured protocols or standardised guidelines, which would enable a more accurate evaluation of the research effort within this field of study.

### Conclusions

Despite limitations such as non-standardised sampling methods and variable publication types and data, this review provides valuable insights into current research trends and gaps regarding the Mediterranean’s marine fauna over the last 130,000 years. Employing the PRISMA method proved effective in synthesising information from different sources, offering a clear framework to identify research gaps. We propose more collaborations between researchers and multidisciplinary areas to develop and improve practical approaches to gathering data in a standardised format for posterior research. This will lead to a broad perspective on marine ecosystem research, along with current and future environmental management research of the Mediterranean Sea’s ecosystems.

## ACKNOWLEDGEMENTS

This work was supported by the Erasmus+ Program Grant no. 2022-1-PT01-KA131-HED-000061155. The author acknowledges the institutional support of the Severo Ochoa Centre of Excellence accreditation (CEX2019-000928-S). This research contributes to the objectives of Q-MARE (a PAGES working group). This research is part of the Integrated Marine Ecosystem Assessments (iMARES) research group funded by Agència de Gestió d’Ajuts Universitaris i de Recerca (Generalitat de Catalunya) Grant no. 2021 SGR 00435.

## DATA AVAILABILITY

All data produced for this work are available in the main manuscript and the supplementary material. The data provided by Leal et al. (2025) regarding the database and timetable will be made public when the current manuscript is published using the same DOI. To access it now use the following link: https://zenodo.org/records/14726420?preview=1&token=eyJhbGciOiJIUzUxMiJ9.eyJpZCI6ImJmNjEyMzQxLWZjMjAtNDUzNy1iZjM0LWU1MGFhMjI0NGMwZiIsImRhdGEiOnt9LCJyYW5kb20iOiJlNTJlMzhmMGFkYjM2Y2MwZDlkOGJkZDBhNWRjOGU0MyJ9.rcMjMNeRD5mMLsCrZLb89VTAoMTQSbN3FDE8IAuXI0t92frryHiEnP23Y3LrN2EOx9RfDi8NUCVonorCl_whJg

## Supplementary Material

### S1. List of Search Strings used in the search engines

**Table S1.** List of Search Strings used in the search engines, organised my theme. Each set of terms divided by theme was combined using the Boolean operator “AND”.

| Topic | Strings |
| --- | --- |
| Taxa | “Delphinus” OR “Monachus” OR “Dolphin*” OR “Porpoise*” OR “Monk seal*” OR “Engraulis” OR “Sardin*” OR “Sparus aurata” OR “Diplodus” OR “Clupea” OR “Alosa” OR “Thynnus” OR “Anchov*” OR “Pilchard*” OR “Bang*” OR “Black sprat*” OR “Fry-dr*” OR “Herring*” OR “*Bream*” OR “Gilt*” OR “Snapper*” OR “Breca*” OR “Puntazzo*” OR “Blacktail*” OR “Tun*” OR “Horse mackerel*” OR “Squid hound*” OR “Ostrea” OR “Hexaplex trunculus” OR “Phorcus turbinatus” OR “Trunculariopsis trunculus” OR “Murex trunculus” OR “Monodonta turbinata” OR “Osilinus turbinatus” OR “Oyster*” OR “Murex*” OR “Turbinate monodont” |
| Area | “Mediterranean” OR “Adriatic” OR “Aegean” OR “Levant*” OR “Tunisia*” OR “Sidra” OR “Ionian” OR “Alboran” OR “Balearic” OR “Tyrrhenian” OR “Ligurian” OR “Sicil*” OR “Libyan” OR “Sardinia” OR “Marmara” OR “Cicilian” OR “Gibraltar” |
| Timeframe | “Holocene” OR “Greenlandian” OR “Northgrippian” OR “Meghalayan” OR “Pleistocene” OR “Preindustrial” OR “Pre-industrial” OR “Pre industrial” OR “Historical” OR “Archaeological” OR “*Lithic” OR “Aurignacian” OR “Gravettian” OR “Solutrean” OR “Magdalenian” OR “Bronze Age” OR “Iron Age” OR “Copper Age” OR “Roman*” OR “Greek*” OR “Medieval” OR “Dark Age*” OR “Little Ice Age” OR “Byzantine” OR “Hellen*” OR “Archaic*” OR “Hunter-Gather*” OR “Hunter* Gather*” OR “Classic*” OR “Antiquity” OR “Christian*” OR “Prehuman” OR “Pre human” OR “Pre-human” OR “Modern” |
| Impacts | “Climat*” OR “Environment*” OR “Salin*” OR “Dissolved oxygen” OR “Productivity” OR “Invasion*” OR “Temperature*” OR “Anthropogenic*” OR “Human*” OR “Whaling” OR “Hunt*” OR “Fish*” OR “Farming” OR “Aquaculture” OR “Harvest*” |
| Theme | “Marine” OR “Maritime” OR “Sea*” OR “Ocean*” OR “Lagoon*” OR “Delta*” OR “Basin*” OR “Estuar*” OR “Shel*” OR “Coast*” OR “Margin*” OR “*Littoral” OR “Gulf*” OR “Strait*” OR “Channel*” OR “Water*” OR “Slope*” OR “Deep” OR “*Tidal” |

### S2. Publications included in the database

**Table S2.**
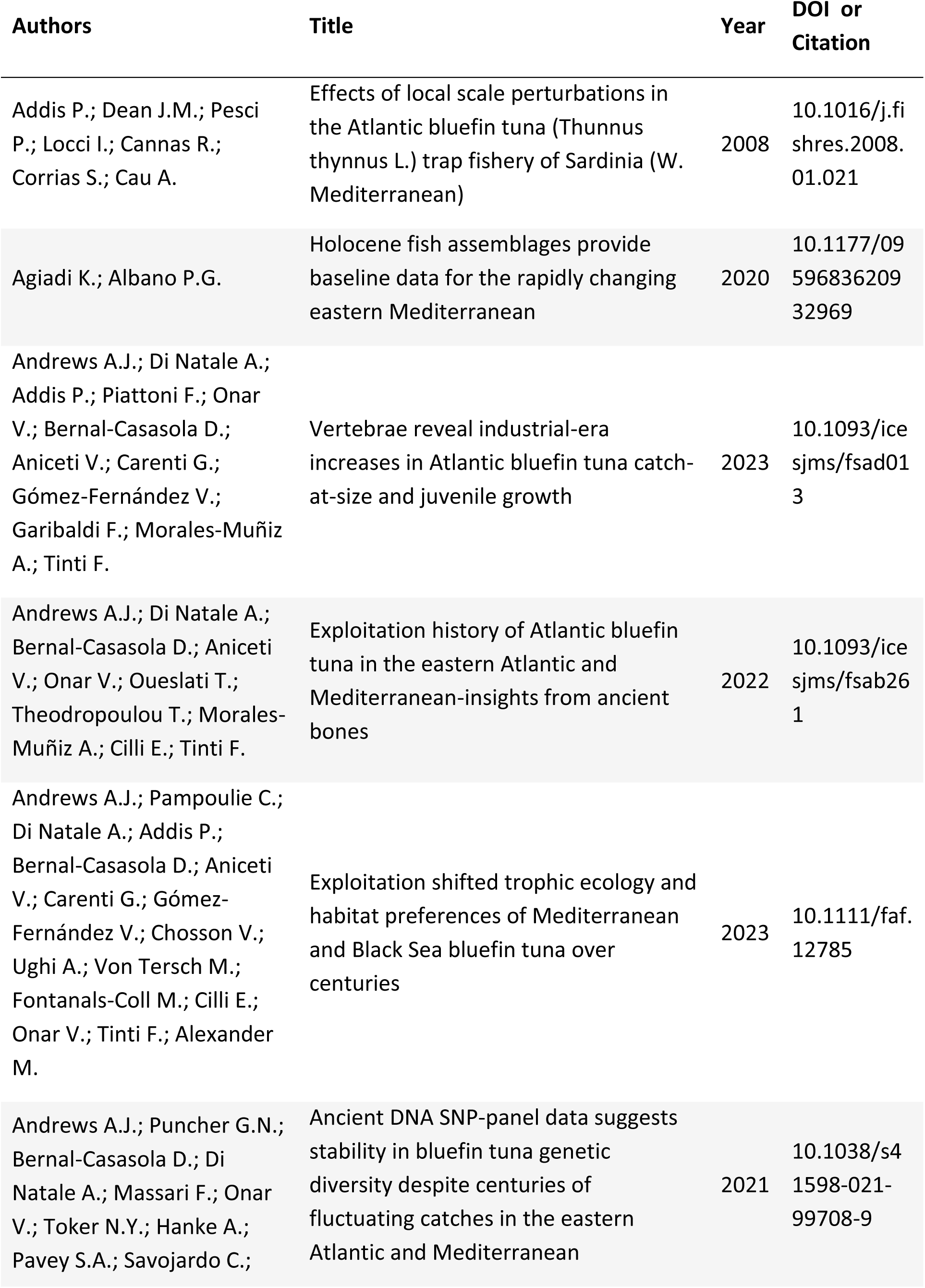
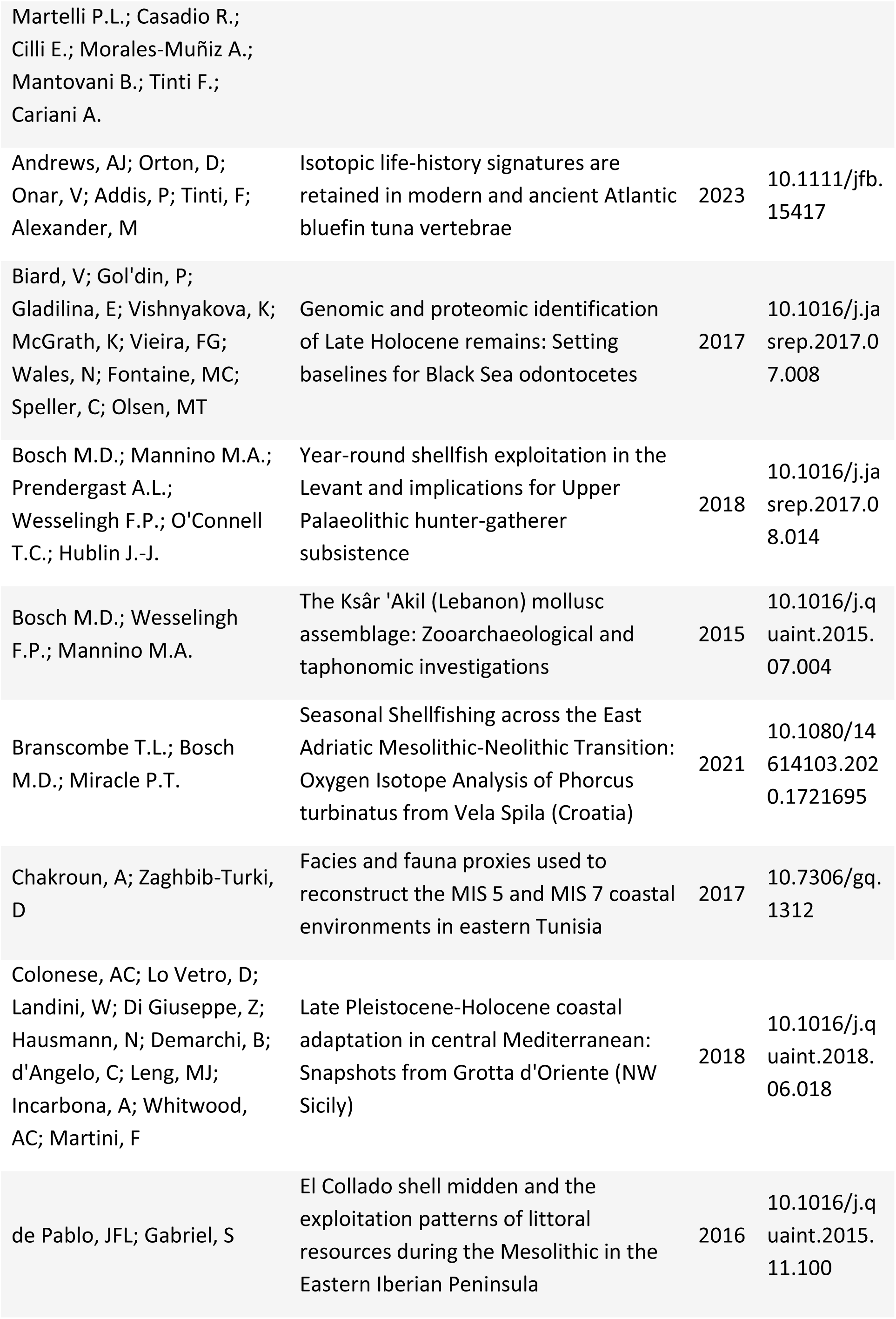
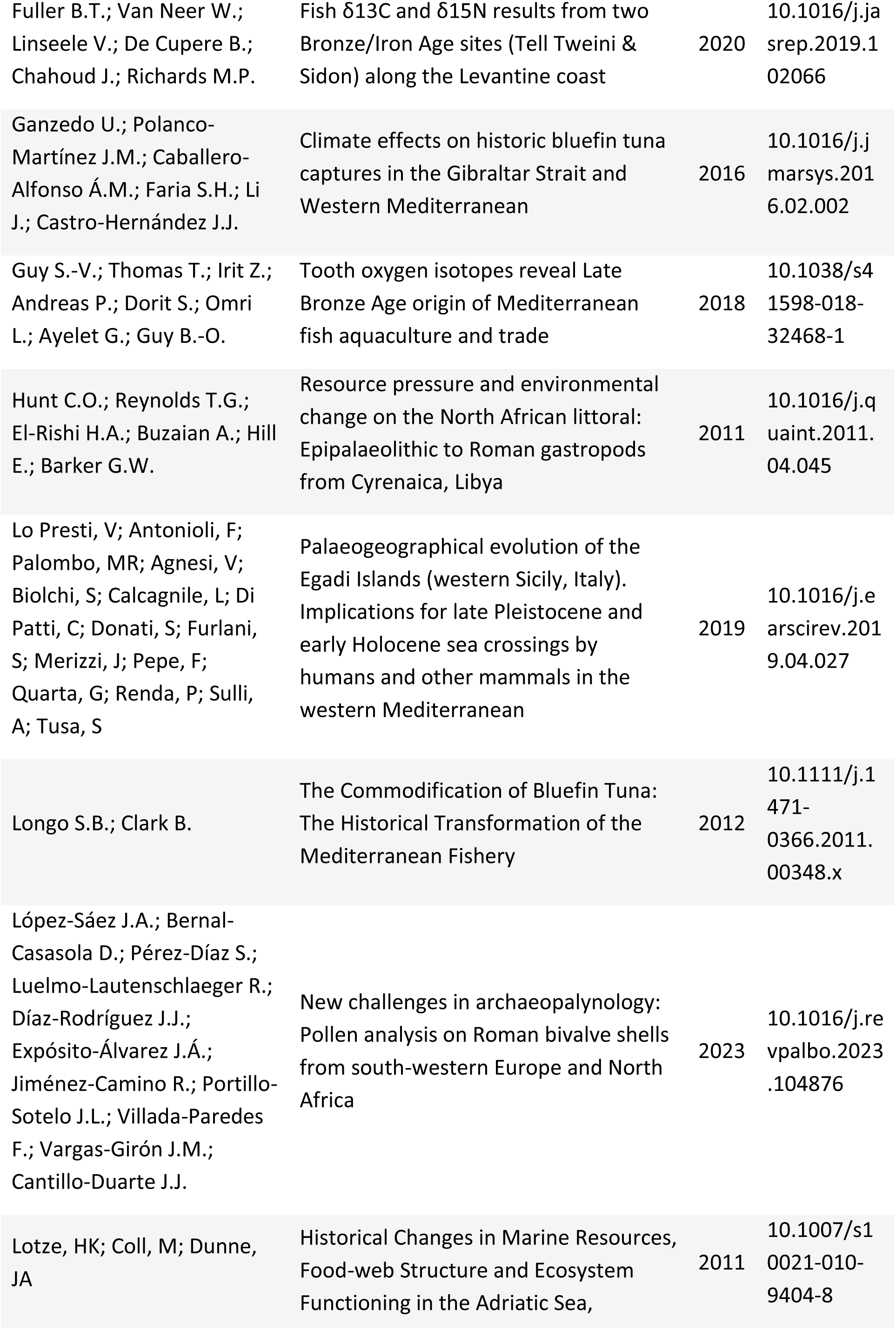
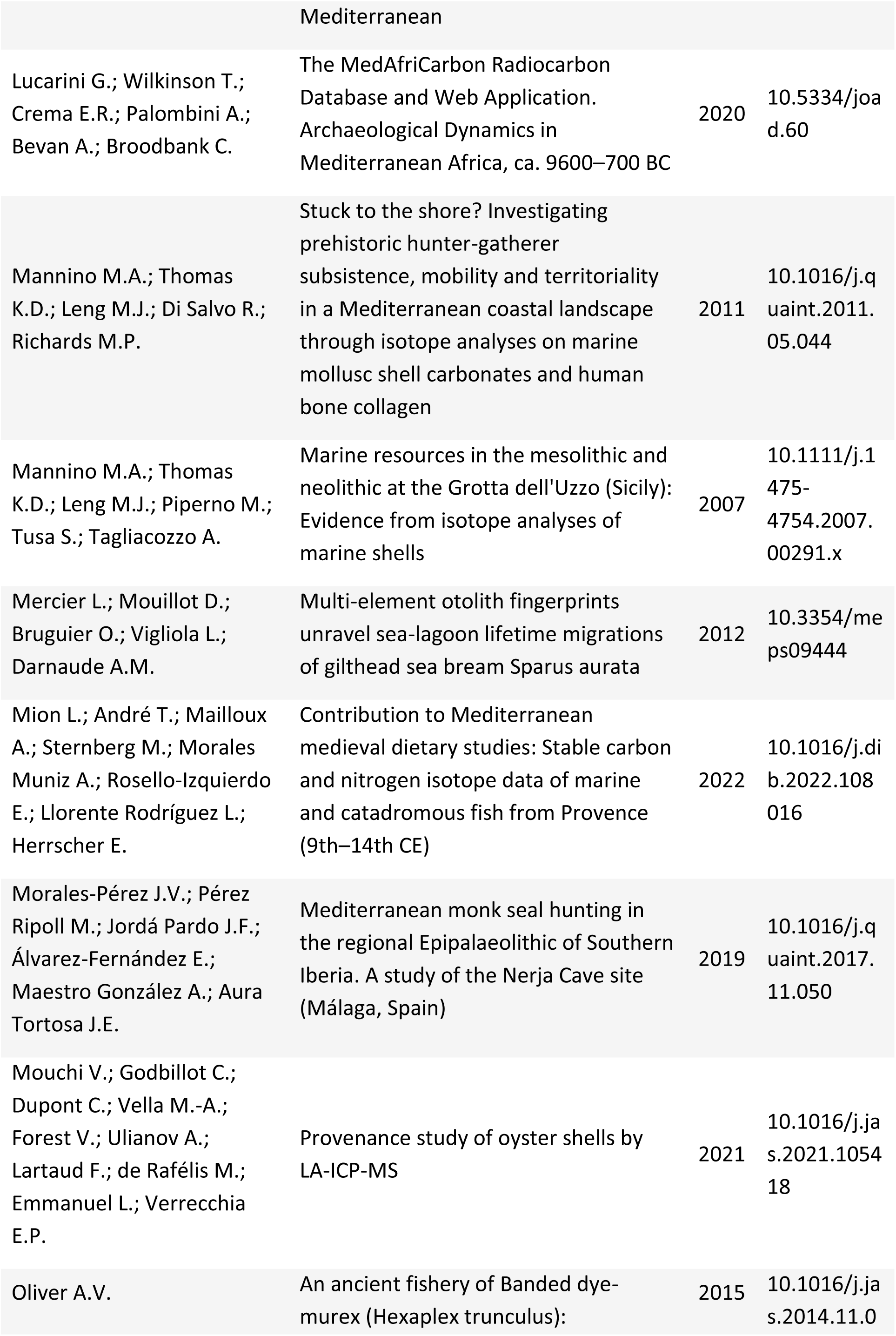
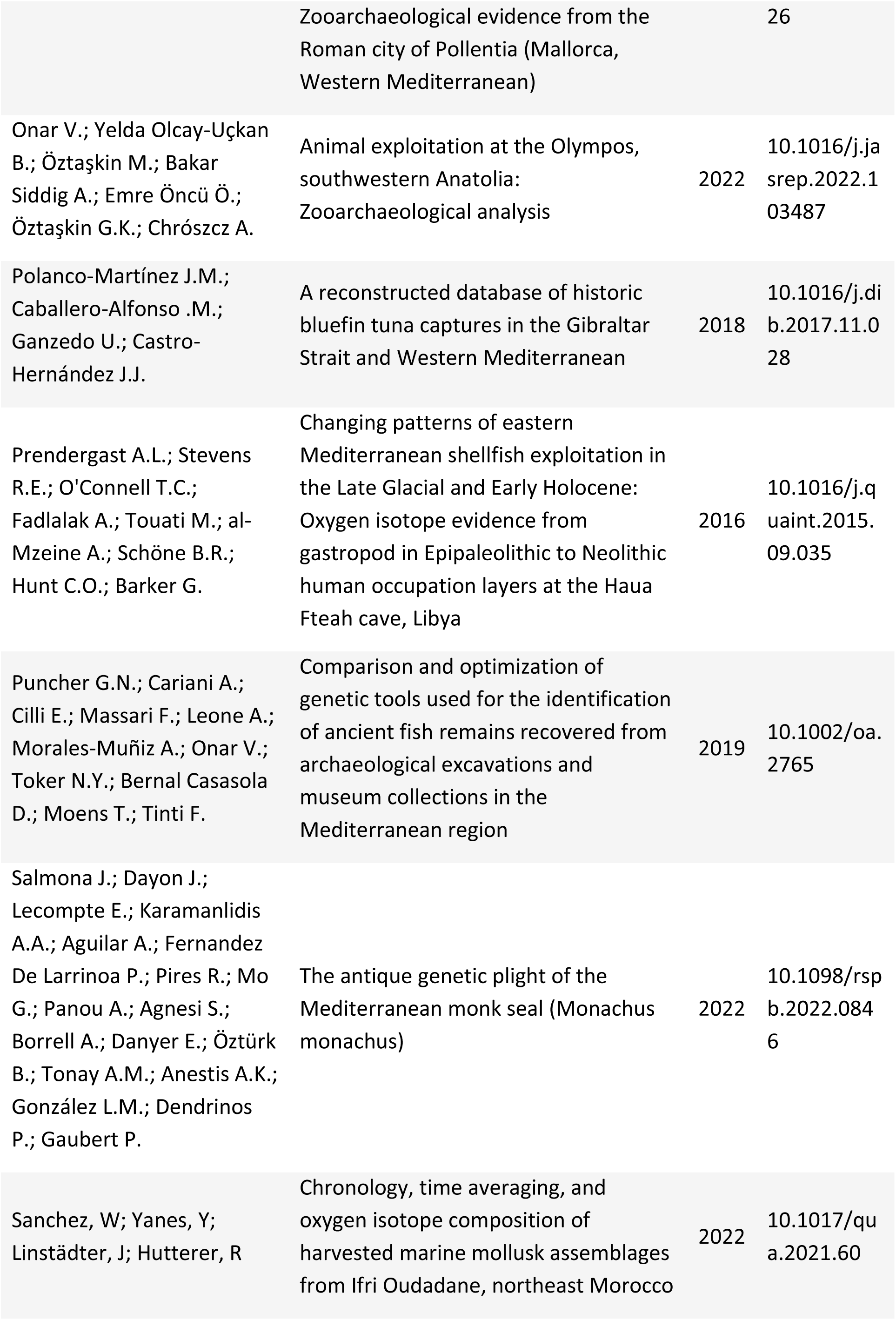
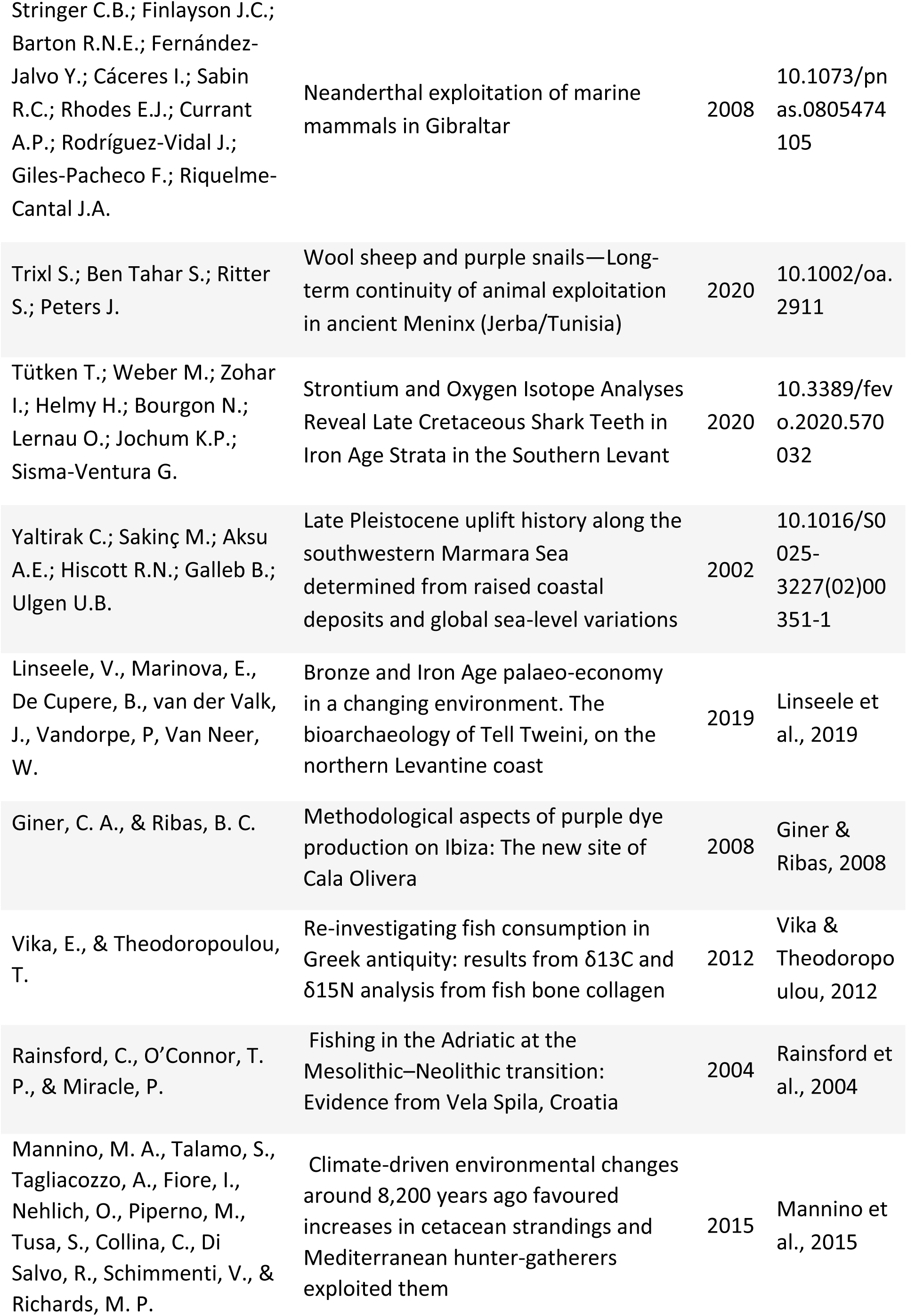
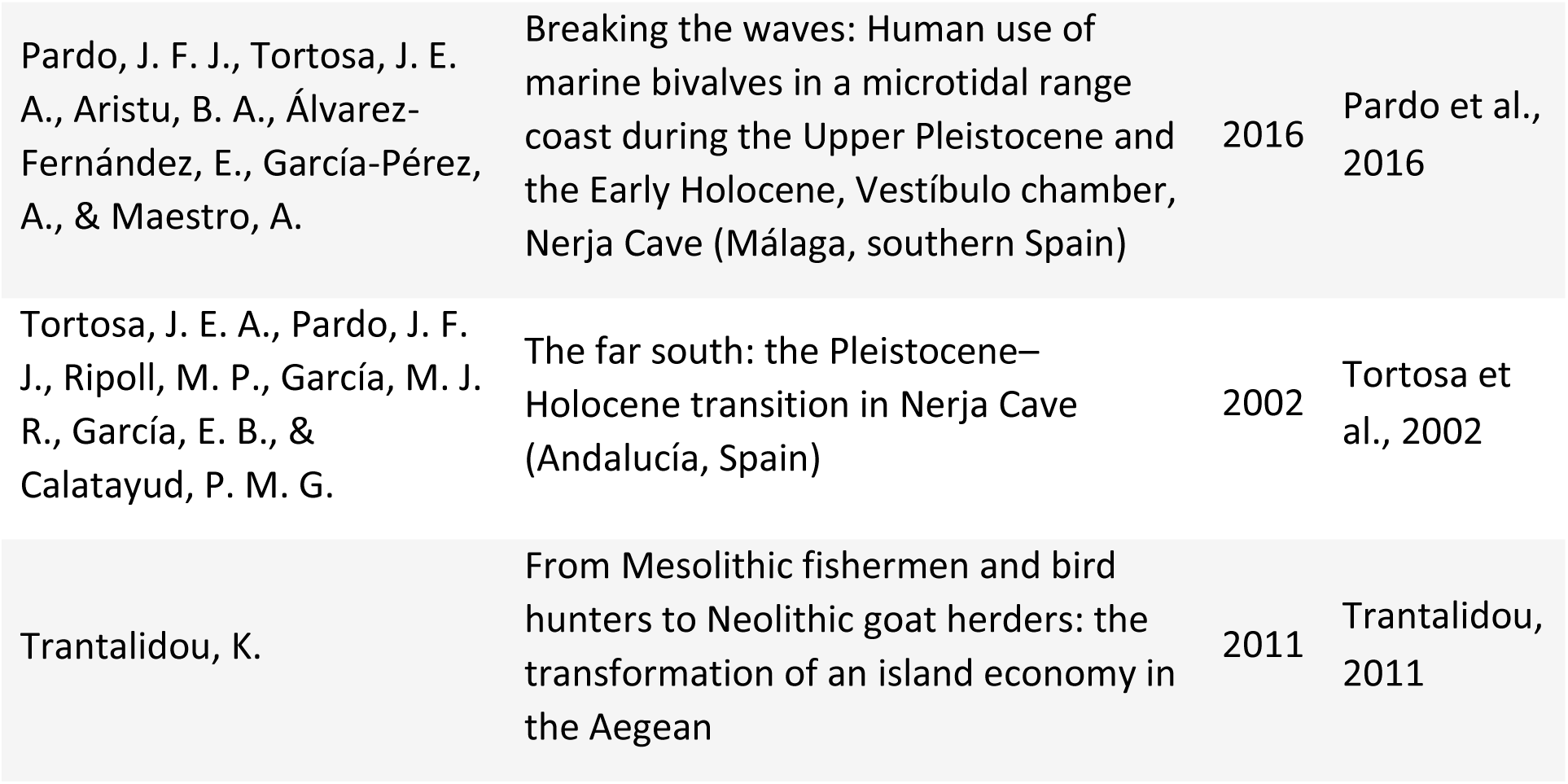
Compilation of the scientific publications used to build the database.

### S3. Table of the Sampling units from the scientific articles

**Table S3.**
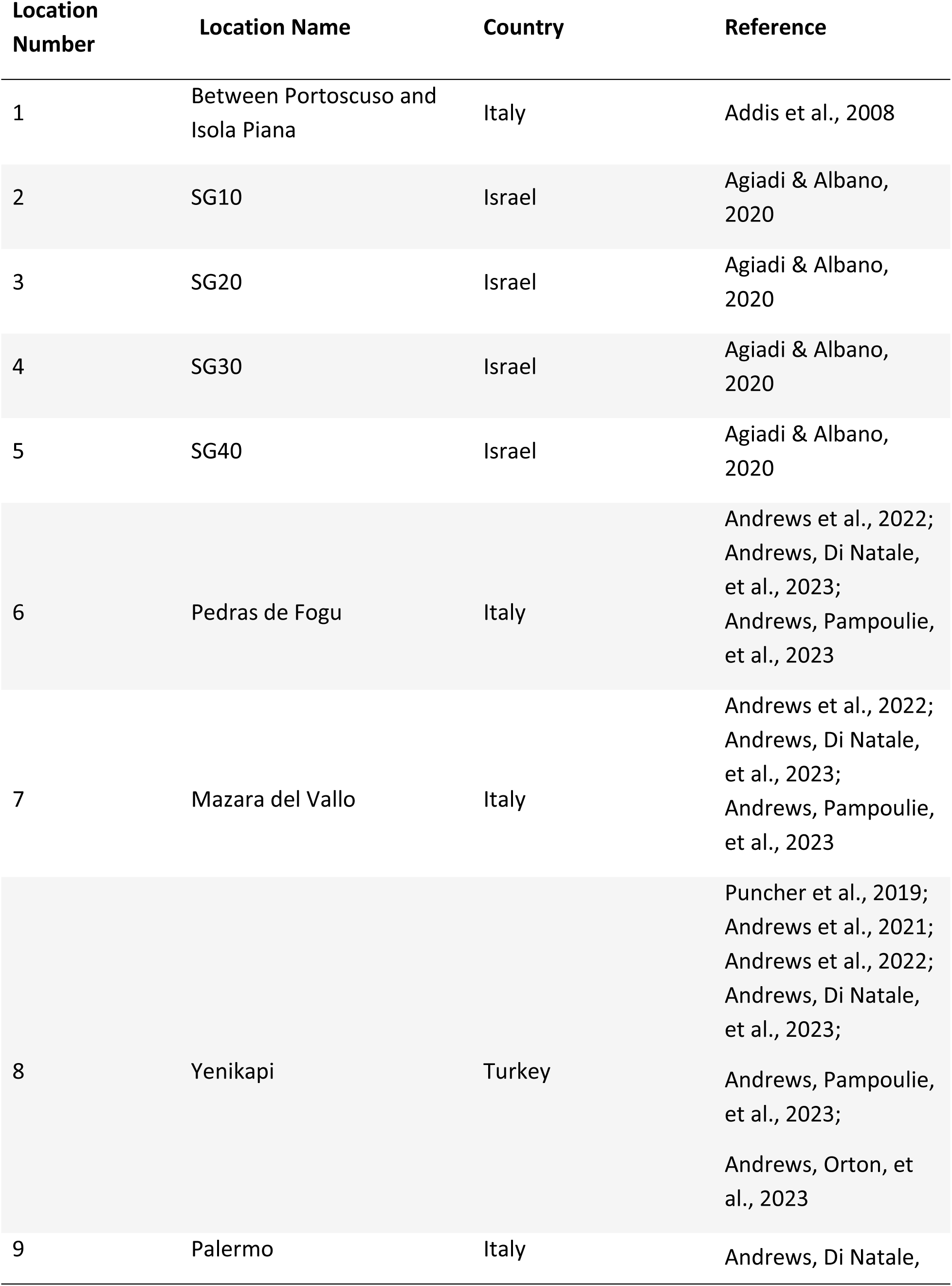
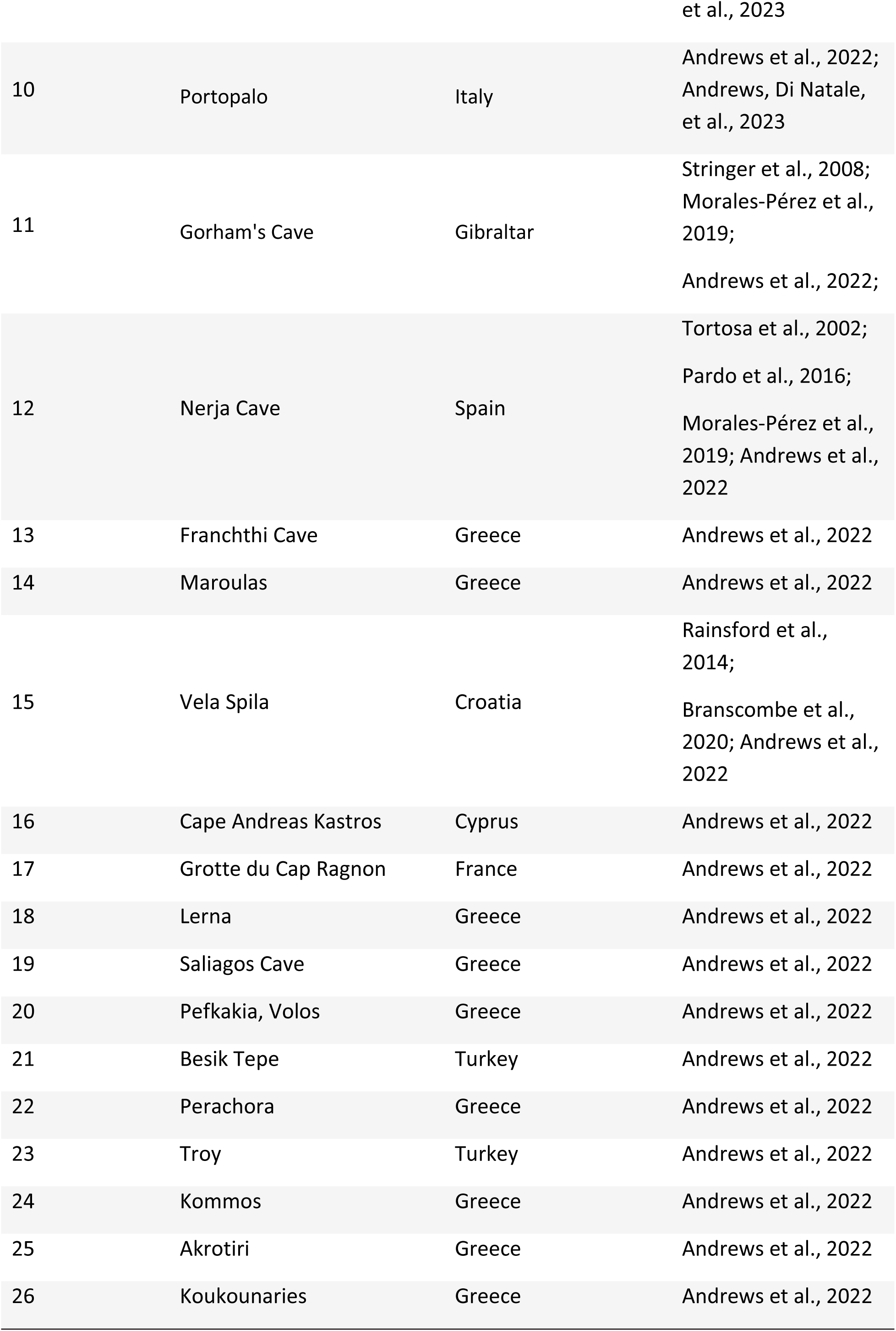
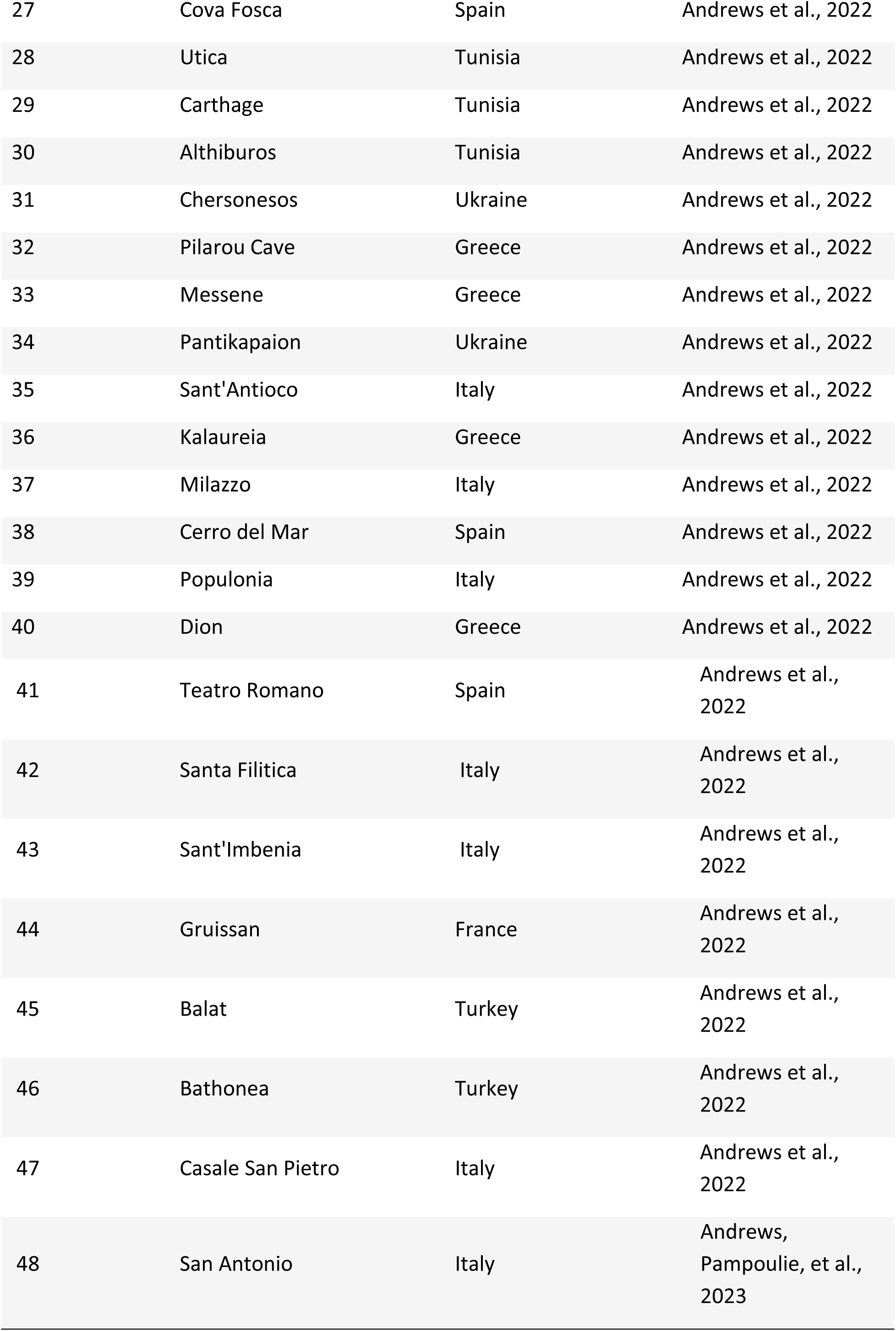
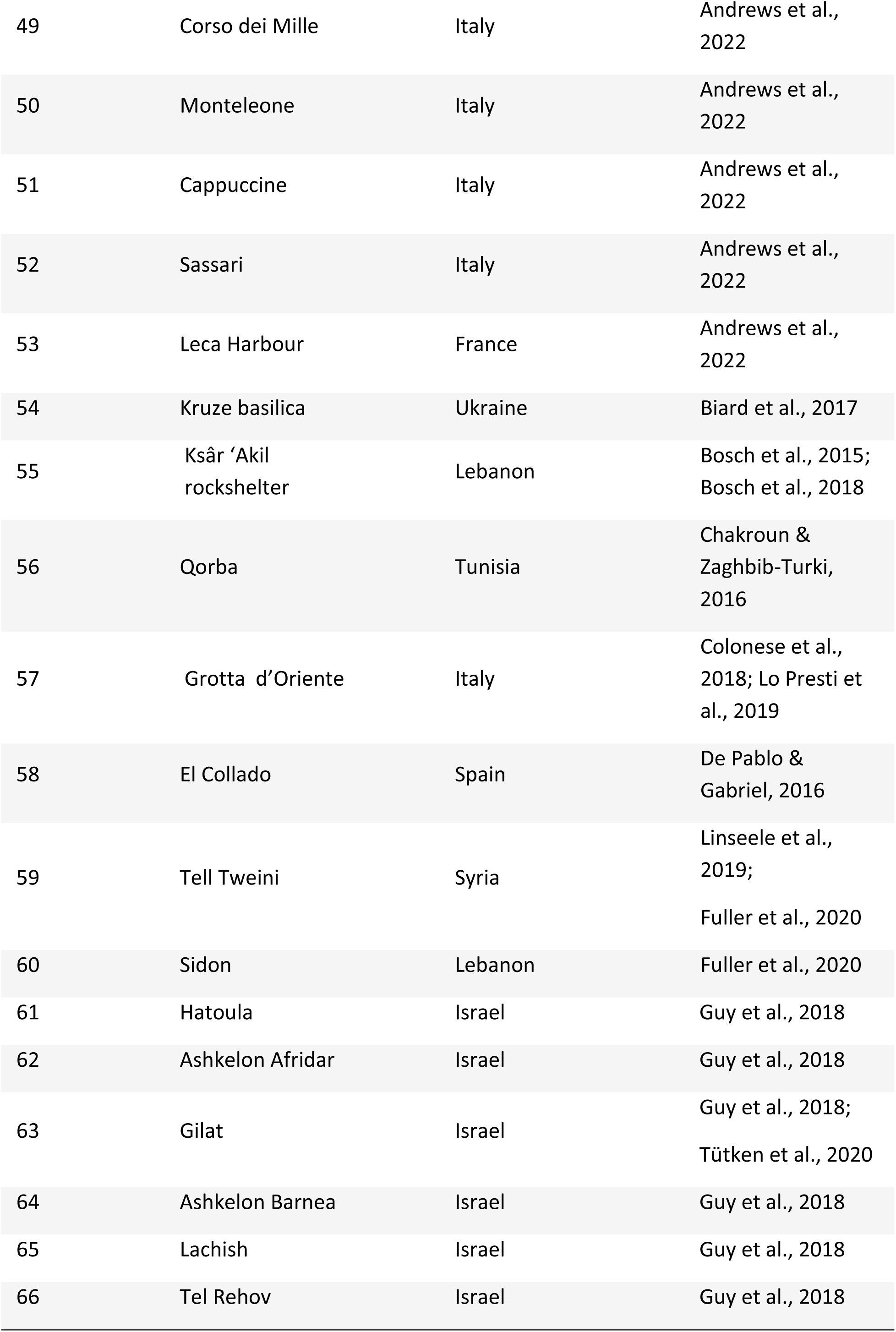
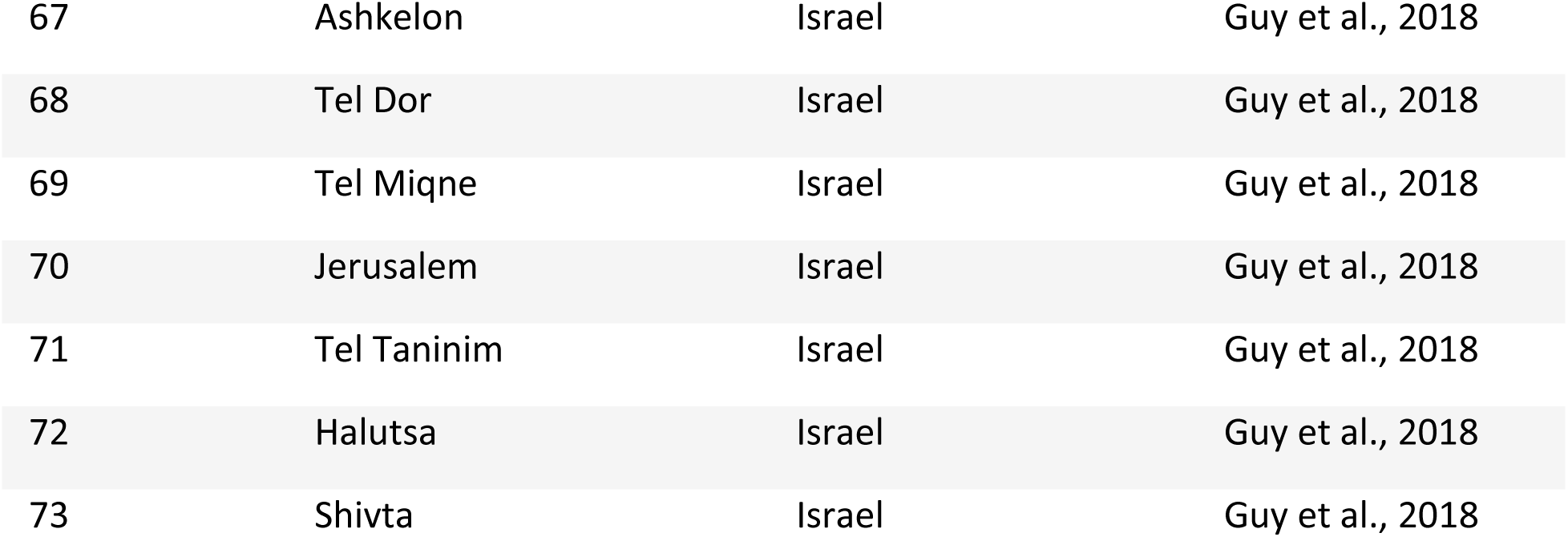
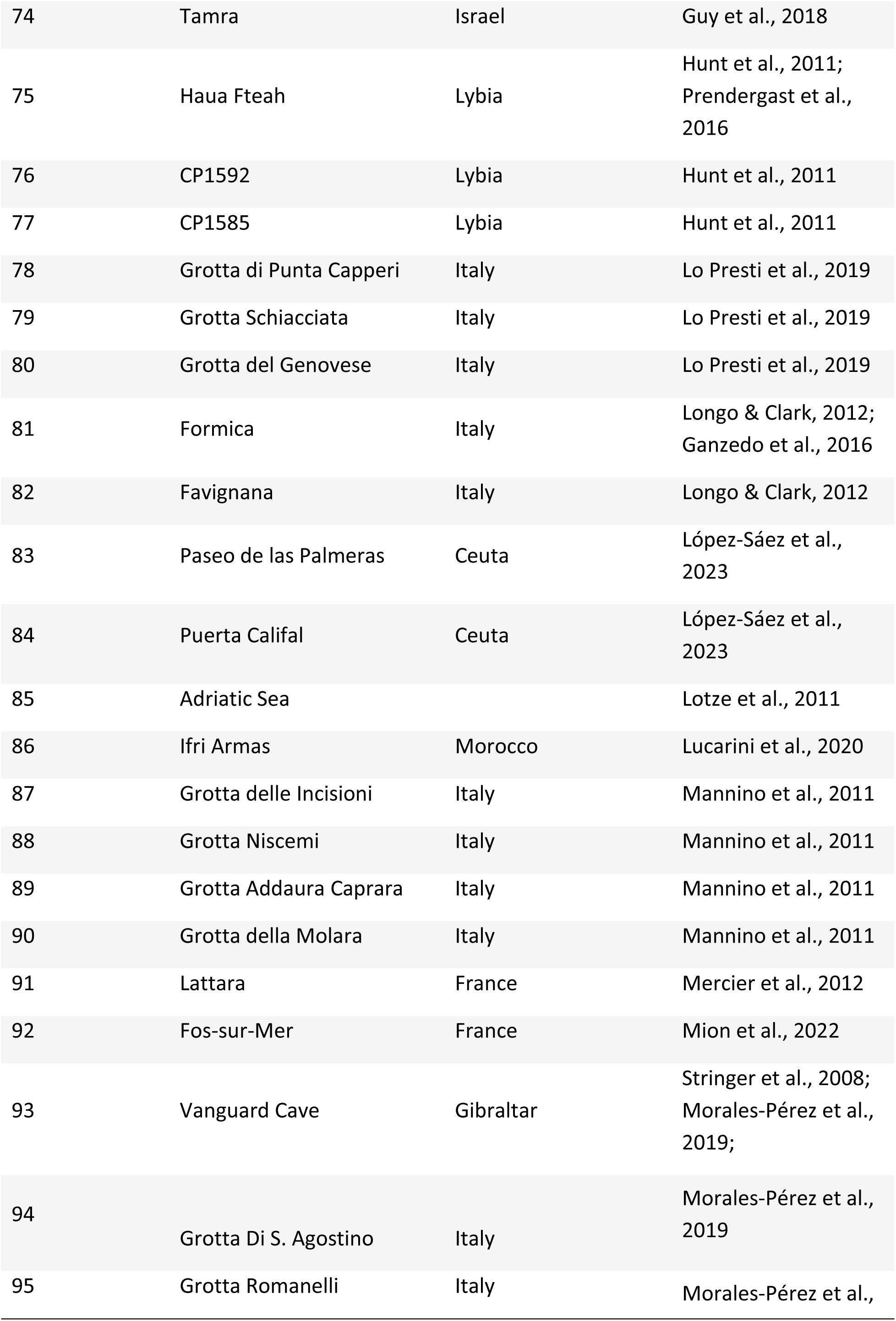
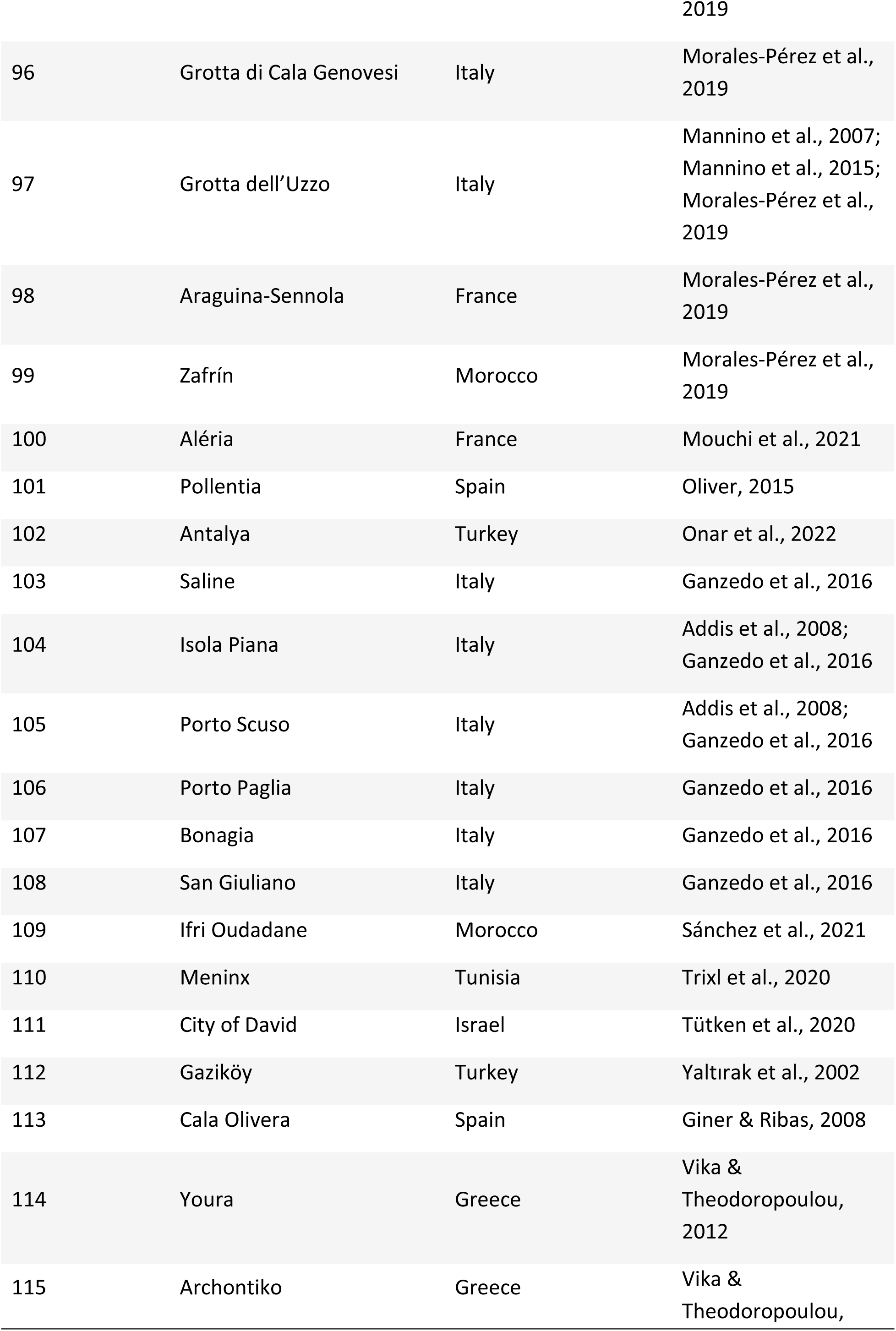
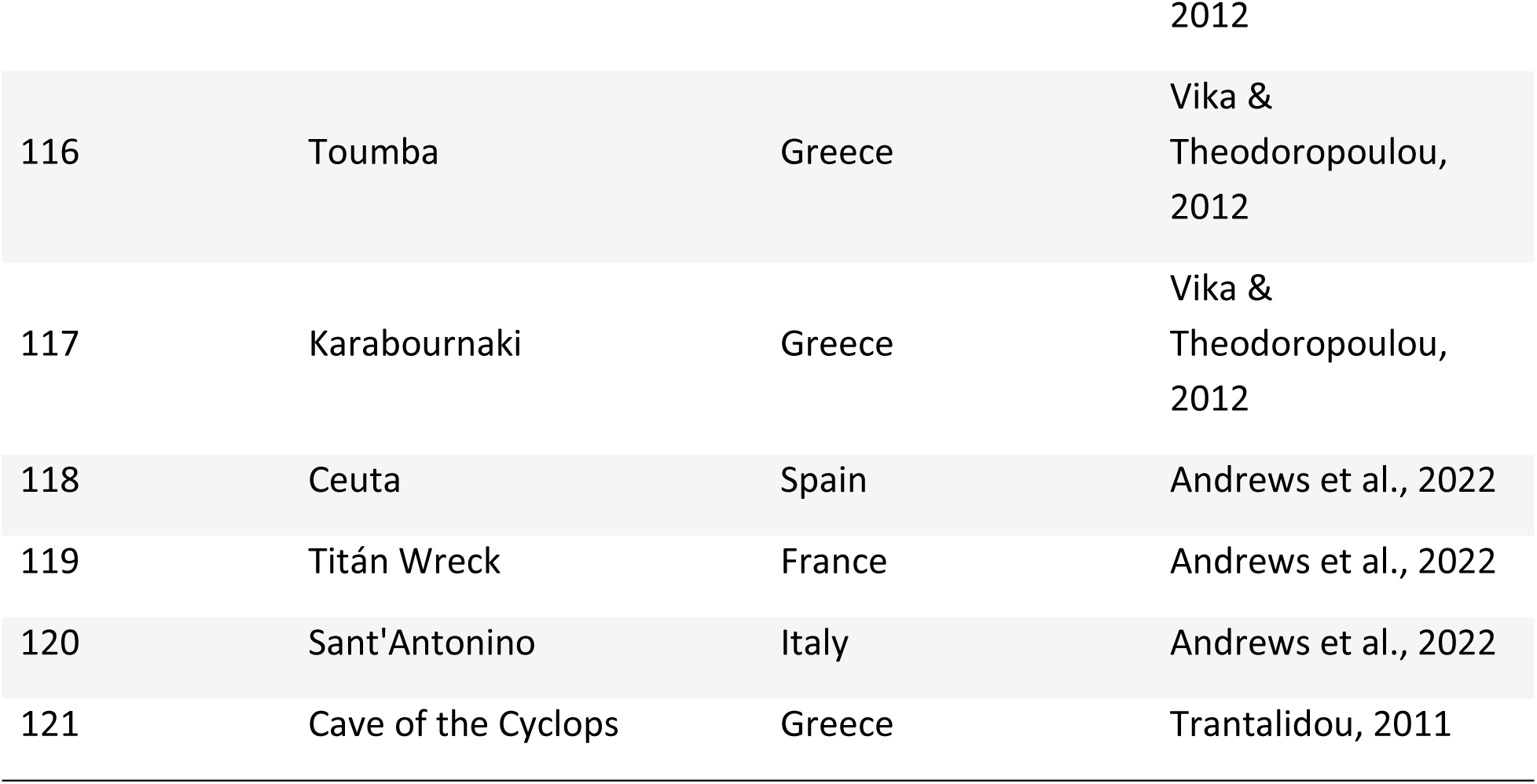
List of location’s number, location’s name, country and respective reference for every sampling unit used in this work.

